# A Parasite Coat Protein Binds Suramin to Confer Drug Resistance

**DOI:** 10.1101/2020.06.04.134106

**Authors:** Johan Zeelen, Monique van Straaten, Joseph Verdi, Alexander Hempelmann, Hamidreza Hashemi, Kathryn Perez, Philip D. Jeffrey, Silvan Hälg, Natalie Wiedemar, Pascal Mäser, F. Nina Papavasiliou, C. Erec Stebbins

## Abstract

Suramin has been a primary early-stage treatment for African trypanosomiasis for nearly one hundred years. Recent studies revealed that trypanosome strains that express the Variant Surface Glycoprotein VSGsur possess heightened resistance to suramin. We show here that VSGsur binds tightly to suramin, other VSGs do not, and that together with VSG13 it defines a structurally divergent subgroup of these coat proteins. The co-crystal structure of VSGsur with suramin reveals that the chemically symmetric drug binds within a large cavity in the VSG homodimer asymmetrically, primarily through contacts of its central benzene rings. Structure-based, loss-of-contact mutations in VSGsur significantly decrease the affinity to suramin and lead to a loss of the resistance phenotype. Altogether, these data show that the resistance phenotype is dependent on the binding of suramin to VSGsur, establishing that the VSG proteins can possess functionality beyond their role in antigenic variation.

## Introduction

Multiple species of the genus *Trypanosoma* cause Sleeping Sickness in humans and related diseases in animals. These vector-borne illnesses are transmitted by the tsetse flies (*Glossina* spp.) and are primarily a burden in sub-Saharan Africa, impacting human populations directly through infection and indirectly through morbidity of livestock^1,2^. There are two subspecies that account for human infections: *T. brucei gambiense* (central and West Africa) and *T. brucei rhodesiense* (east and southern Africa). Numerous animal-pathogenic subspecies exist causing nagana in cattle (*T. brucei brucei, T. congolense, T. vivax*), surra in camels and horses (*T. evansi, T. equiperdum*), and trypanosomiasis in pigs and wild boars (*T. simiae*)^3^.

A remarkable feature characterizing all such infections is the ability of the trypanosomes to thrive in the blood and tissue spaces of infected mammals despite complete exposure to the immune system of the host^4^. This is achieved largely through a dedicated machinery in the organism to alter its surface coat and continuously evade immune system recognition and clearance^5^. The trypanosome surface is densely carpeted with approximately 10 million copies of the Variant Surface Glycoprotein (VSG), a GPI-anchored and glycosylated polypeptide to which the immune system mounts a very effective response^6,7^. The VSGs are long, rod-shaped homodimeric proteins of around 60kDa with two subdomains: a larger, N-terminal domain (NTD) of around 350-400 amino acids and a smaller 80-120 residue C-terminal domain (CTD) to which the GPI anchor is attached (Supplementary Fig. 1)^8^. The NTDs consist of a conserved three-helix bundle scaffolding, the loops connecting the helices adorned with large insertions forming top and bottom subdomains or lobes (Supplementary Fig. 1). The top lobe (facing away from the parasite) is hypothesized to harbor the majority of immune epitopes, presenting the “antigenic face” of the VSG, although little epitope mapping and no antibody-VSG co-crystal structures have been published to date^8–10^.

While the host immune system develops rapid and effective responses to the VSGs, the trypanosomes access a genetic repository of over 2000 VSG genes and pseudo-VSG genes with distinct antigenic properties, switching coat proteins and thereby rendering the host response naïve to the new coat before the parasites can be cleared. This process of recognition, partial clearance, coat switching, and pathogen escape is termed antigenic variation. Because of this ability to thwart immune clearance, African trypanosomiasis is usually fatal unless treated chemotherapeutically^4^.

Synthesized as early as 1917, the compound suramin (originally Bayer 205 and later sold as Germanin) was first used therapeutically to treat African trypanosomiasis in the 1920s^11–13^. It is one of only a few drugs (including pentamidine, melarsoprol, eflornithine, nifirtimox, and fexinidazole)^12,13^ available to counter the disease and is only effective in the early stages of infection, before the parasite has entered the central nervous system (as suramin cannot effectively penetrate the blood-brain barrier^14^). It has also been used prophylactically^15^ and is on the World Health Organization’s List of Essential Medicines^16^. Recently, suramin has demonstrated potent antineoplastic properties^17^. It is part of a larger family of benzopurpurine dyes and naphthalene ureas that show trypanocidal activity^12^. Resistance to suramin has gradually spread through the animal-infective trypanosomal populations, but has yet to reach those species causing disease in humans^18^.

The specific mechanism behind the trypanolytic activity of suramin remains unresolved, although studies have implicated effects on the glycosome and impairment of cytokinesis^19,20^. A model for the internalization of suramin has been proposed in which the drug enters through receptor-mediated endocytosis involving two distinct pathways: (1) low-density lipoproteins (LDLs) with possible involvement of other serum proteins and (2) the invariant surface glycoprotein 75 (ISG75) receptor of the pathogen^21–23^. However, to date no direct binding of suramin to any trypanosomal protein has been reported.

Recently, *in vitro* selection in the presence of suramin generated trypanosome strains that were over ninety-fold resistant to the drug^18^. All these resistant strains were shown to express a specific VSG, termed VSGsur, that itself was sufficient to convey the resistant phenotype in trypanosomes genetically engineered to express it^23^. However, it has remained unclear how VSGsur is involved in this resistance phenotype, whether directly or indirectly. Hypotheses include a model in which VSGsur causes suramin resistance by decreasing specific, receptor-mediated endocytosis pathways critical to the uptake of the drug^23^.

We show here that VSGsur directly binds suramin with high affinity, whereas other VSGs do not. Crystal structures of VSGsur and VSG13 identify a subfamily of VSGs that structurally diverge from all those resolved to date in several aspects, including a cavity between the monomers in the VSG-homodimer interface. The co-crystal structure of VSGsur with suramin shows that the drug binds in this cavity, while a structure-based, single loss-of-contact mutation abrogates both suramin binding and resistance.

## Results

### Structures of VSGsur and VSG13 reveal a divergent subfamily of VSGs

We sought insight into the function of VSGsur through structural studies of the protein (clone from *T. brucei rhodesiense*, Genbank ATI14856), determining the crystal structure of the NTD to 1.2Å resolution (residues 30-408, Fig. 1a, Methods, Supplementary Fig. 2 and Table 1). VSGsur differs markedly from previous VSG structures determined (VSG1, VSG2, VSG3, and ILTat1.24)^24–27^. While the core 3-helix bundle scaffolding of the VSG family is present, VSGsur possesses several divergent features. The most striking is observed in the “upper lobe” of the molecule. The insertion between the helices at the top of VSGsur folds into a *bona fide* structural subdomain (residues ∼140-260) consisting of five β-strands and an elongated loop between 174-208 that travels half the length of the VSG molecule before returning to the upper lobe (Supplementary Fig. 3a). This subdomain forms an intermolecular β-sandwich with the pairing monomer in the homodimer to create a tightly interwoven quaternary fold with a significant hydrophobic core. Perched on top of the 3-helix bundle, this “head” is strikingly different than the more flattened polypeptide arrangements of the other VSG top lobes that do not organize into any typified fold.

**Fig. 1:**
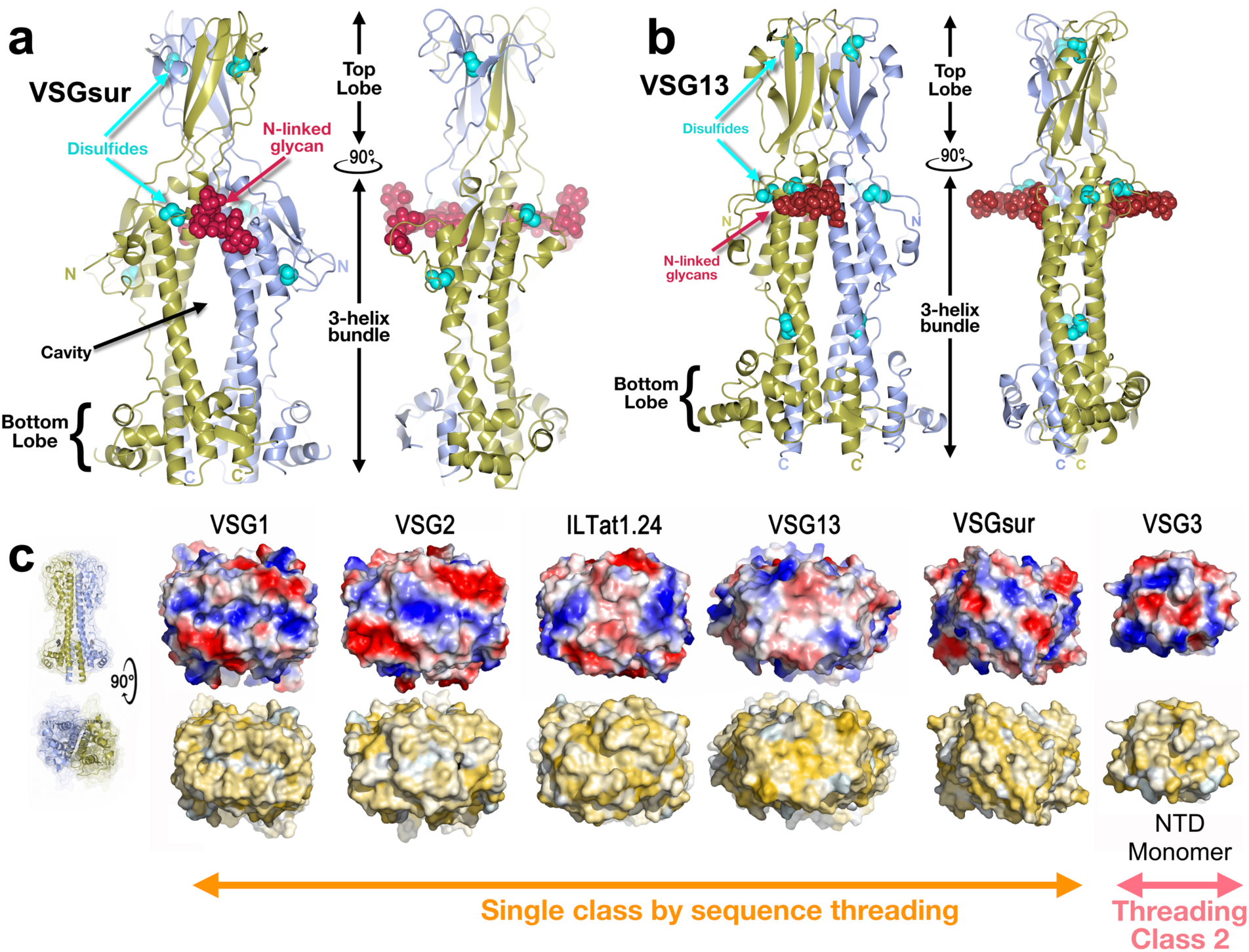
Overall Structure of VSGsur. **(a)** VSGsur is shown as a ribbon diagram in two orientations related by a 90 degree rotation, the monomers in the homodimer colored gold and light blue. The N-linked glycans of the two VSG monomers are displayed as red space-filling atoms, disulfide bonds are shown in cyan, and several key features are labeled (top and bottom lobes, 3-helix bundle, and the cavity between the dimers. “N” and “C” indicate the N- and C-termini of each monomer, color coded to match the ribbon diagrams. **(b)** The crystal structure of VSG13 in two orientations is depicted similarly to that of VSGsur as in (a). **(c)** Molecular surfaces of published VSG structures to date. The VSGs are oriented looking “down” on the top lobe of the protein, the Fig. rotated 90 degrees about a horizontal axis in comparison to previous Fig.s. The upper row of surfaces is colored by relative electrostatic potential (blue is basic/positively charged, red is acidic/negatively charged, and white is neutral) and on the lower row by the Eisenberg hydrophobicity scale^48^ (yellow indicates hydrophobic, white polar). Arrows underneath the surfaces illustrate the two broad superfamilies of different VSGs identified by sequence threading as implemented by PHYRE2^49^. Note that the NTD of VSG3 is monomer in solution and the crystal structure, and in the absence of any structural data of a NTD dimer, we have chosen to show only the monomeric surface. Structures and molecular surfaces illustrated with CCP4mg^50^ and MacPyMOL^51^.

Also remarkable is the location of an N-linked sugar nearly two-thirds of the distance up the NTD from the bottom lobe and directly under the β-sandwich top lobe. Most VSGs possess N-linked carbohydrates, but in all studied to date they are located in the bottom lobe and have been hypothesized to function there in VSG-membrane dynamics^28^. Further distinguishing VSGsur from previous structures, the multiple disulfide bonds stabilizing most VSG proteins (three pairs in VSGsur) are not clustered in the upper lobe, but extend into the 3-helix bundle. Finally, there is a large cavity in the homodimeric interface in VSGsur not found in other published VSG structures, located just beneath the unusually placed N-linked glycan. Altogether, VSGsur is a significant departure from previously studied VSGs while still maintaining the core scaffolding of this family.

Our additional studies show that VSGsur is not a unique fold but is in fact likely a member of a subfamily of VSGs. The 1.4Å resolution crystal structure of VSG13 (also named MITat1.13) reveals a second VSG protein with a top “head” consisting of a large β-sheet subdomain (residues 130-260) that forms an intermolecular β-sandwich in the homodimer (Fig. 1b, Methods, Supplementary Fig. 3 and Table 2). Even more striking than VSGsur, the cysteine disulfide bonds of VSG13 (four in this case) are spread throughout the length of the VSG and not clustered in the top lobe, in fact reaching down near the bottom lobe of the NTD. As with VSGsur, VSG13 harbors an N-linked glycan approaching the top of the molecule, positioned just under the β-sheet head and not in the bottom lobe. VSG13 also possesses an internal cavity between monomers in the homodimer. While smaller than that observed in VSGsur, it is still large enough to allow solvent into the cavity from the crystallization reagents. A protein structural alignment of the VSGsur and VSG13 monomers (Supplementary Fig. 3b) shows that the core 3-helix bundle aligns well and the N-linked sugars superpose very closely, whereas the β-sheet head in the upper lobes align poorly, evincing significant divergence. Comparing VSGsur and VSG13 with the “canonical” VSG2 structure shows that the β-sandwich subdomain gives more height to the NTDs of this subfamily than the other VSGs previously studied (Supplementary Fig. 3c), which could have significance for membrane dynamics and receptor-mediated endocytosis^28^.

Although VSG13 and VSGsur present marked divergences from previously reported VSG structures, the variability in these two Variant Surface Glycoproteins is housed within more generally conserved features that make them clear members of a protein superfamily. Most obvious is the core 3-helix bundle scaffolding that undergirds all the VSGs and VSG-related proteins such as Serum Resistance-Associated protein (SRA)^29^ and the haptoglobin–hemoglobin receptor^30^. The divergent structures in the top and bottom lobes of the VSGs are basically different insertions between the loops connecting the central helices of the core of these rod-like molecules. Secondly, the proteins all evince a shared functionality in antigenic variation by presenting distinct molecular surfaces to the extracellular environment with extensive variation in amino acid sequence, charge and hydrophobic distributions, and topography (Fig. 1c).

### Resistance studies with suramin

To examine whether the resistance mechanism associated with the expression of VSGsur involved direct protein-to-drug binding, we tested *T. brucei rhodesiense* VSGsur (expressed in *T. brucei brucei*) and a panel of other VSGs from *T. brucei brucei* Lister 427 (including VSG13) for their effect on suramin susceptibility and suramin binding (Fig. 2). Toxicity experiments demonstrated that, similarly to what has been shown for several VSGs from *T. brucei rhodesiense* and *T. brucei brucei* strain BS221^18,23^, suramin killed all VSG expressing Lister 427 strains we examined (VSG2, VSG3, and VSG13), whereas those expressing VSGsur were resistant to 25-40x higher concentrations of the drug (Fig. 2a), while showing the same growth rate in the presence of 0.7 µM suramin as in the absence of the drug (doubling time 6.5 h, Supplementary Fig. 4). Isothermal titration calorimetry (ITC) showed that while suramin did not bind to VSG2, VSG3, or VSG13, it bound with nanomolar affinity to VSGsur (Fig. 2b and Supplementary Fig. 5).

**Fig. 2:**
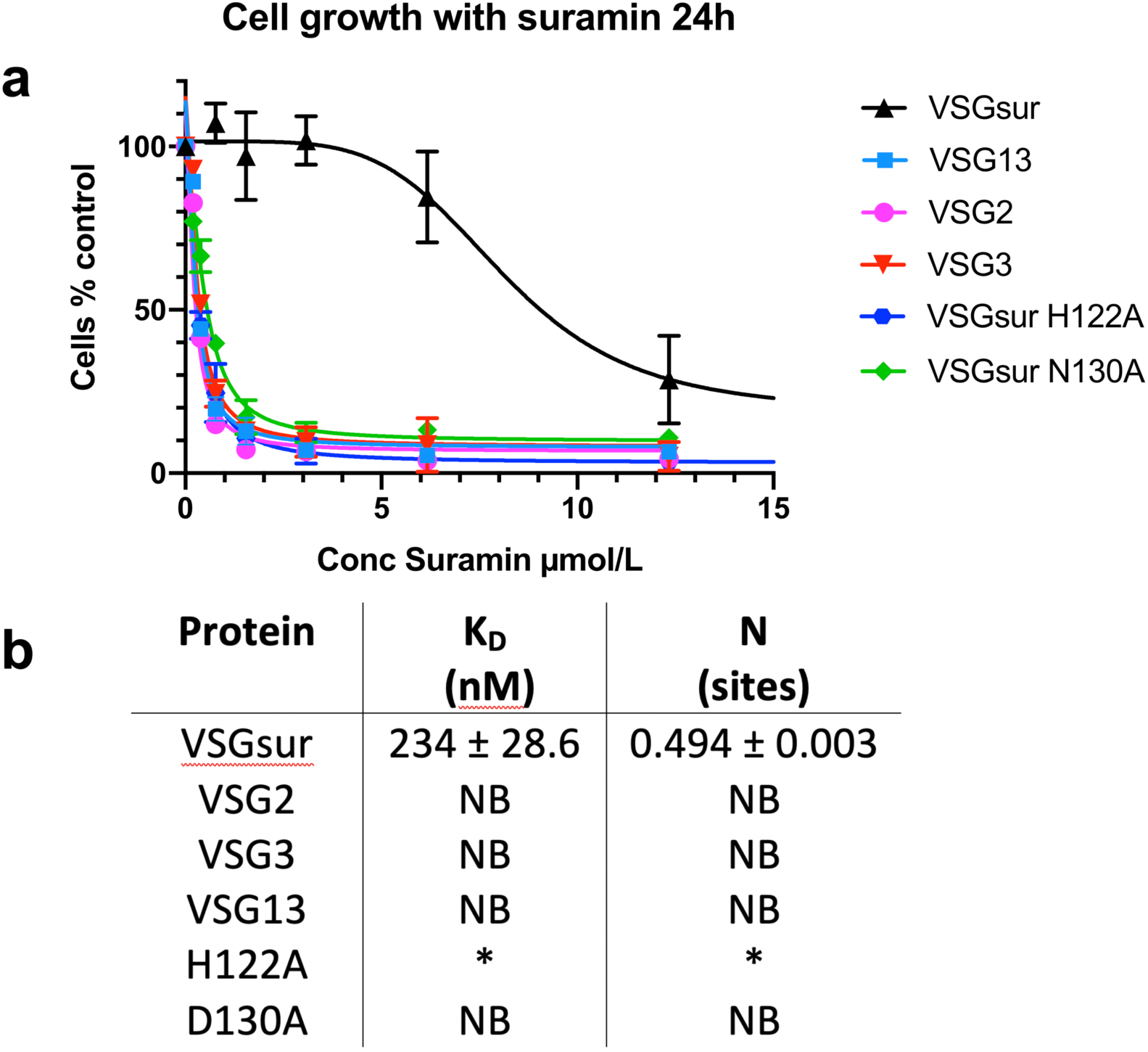
Suramin Binds only VSGsur and is toxic to all other VSGs examined. **(a)** Growth curves at different suramin concentrations show that VSG2, VSG3, and VSG13 are sensitive to the drug and only VSGsur shows resistance, whereas two structure-based mutants in VSGsur (H122A and N130A) lose this resistance. (n = 3 sampled). **(b)** Isothermal calorimetric results of suramin binding to different VSGs. Matching the toxicity assays, suramin was found to bind tightly only to wild type VSGsur, but not well to the two structure-based mutants (H122A and N130A) or VSG2, VSG3, and VSG13. “NB” indicates no binding. “*” for the VSGsur H122A mutant indicates evidence of weak binding (Supplementary Fig. 5), but a satisfactory fit to the curve was not possible. The binding for H122A is expected to be in the high µM range.

### Co-crystal structure of VSGsur with suramin

The above data motivated efforts to co-crystallize suramin together with VSGsur. Soaking native crystals in high concentrations of suramin led to well-diffracting crystals into which the drug could be modeled (Methods, Supplementary Table 1). Interestingly, the chemically symmetric drug suramin bound at the dimerization interface *asymmetrically* (Fig. 3a). Suramin binds tightly in the large cavity between the two monomers of VSGsur, burying a surface area of approximately 700Å^2^ with more than one hundred atomic contacts characterized by both hydrogen bonding and non-bonding interactions. Suramin (C_51_H_40_N_6_O_23_S_6_) is a symmetric, polysulphonated naphthylurea (Supplementary Fig. 6a). The compound is centered on the urea functional group (NH–CO–NH). Extending from the urea moiety in each direction are a pair of benzene rings (thus a total of four in the molecule), each connected by an amide and finally linking to a terminal naphthalene decorated with three sulfonic acid groups (therefore, two naphthalenes and six sulfonic acids in total, Supplementary Fig. 6a). The rotation axis of the VSGsur dimer passes through one of the benzene rings connected to the urea group (Fig. 3a and Supplementary Fig. 6b). Thus, the central urea group of suramin binds just “off center” with respect to the dimer rotation axis. Three of the four benzene groups in the center of the suramin molecule bind in the middle of the dimer interface. These are well-ordered in the structure, with clear electron density and B-factors lower than the average of the overall structure (protein, ligands, and solvent). The naphthalene rings, however, show weaker electron density and high B-factors, characteristic of more disorder or conformational flexibility. Their extension outside the cavity results in fewer contacts from VSGsur to stabilize their conformation.

**Fig. 3:**
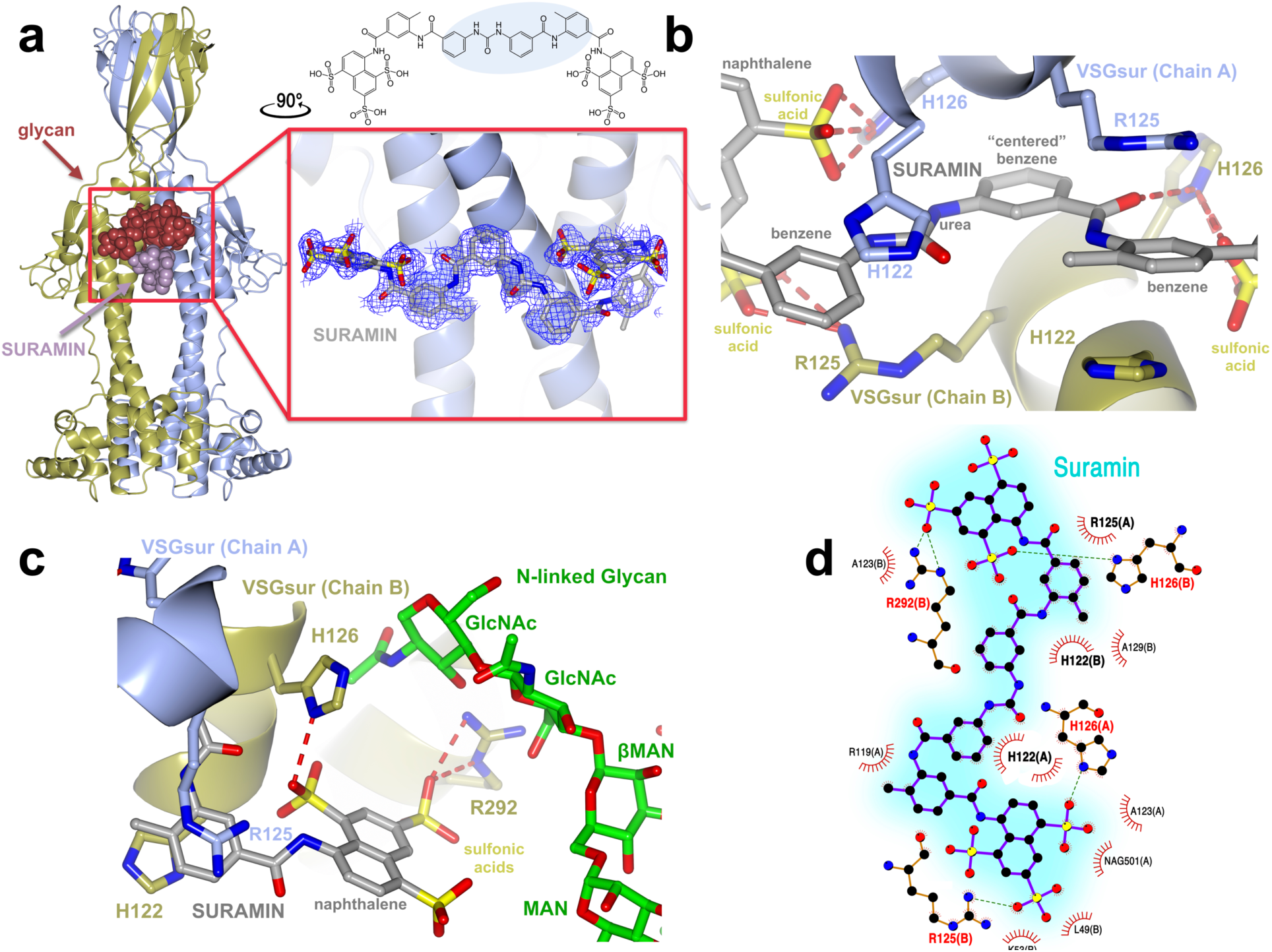
Suramin binds in the cavity between the VSGsur monomers. **(a)** On the left is the overall structure of the VSGsur-suramin co-crystal structure in ribbon diagram form similar to that depicted in Fig. 1. Suramin is shown in purple as a space-filling representation. The red box around the drug is expanded in the boxed image on the right (rotated 90 degrees counterclockwise about a vertical axis), showing a ball-and-stick rendering of the compound with a blue wire-cage of the 1σ contour of the 1.9Å 2Fo-Fc electron density map of the refined molecule drawn around it. Suramin is shown as a ball-and-stick chemical representation with oxygen atoms are shown in red, nitrogen in blue, and carbon gray. **(b)** VSGsur is depicted as in Fig. 1 and suramin as in panel (a). Side-chains from VSGsur are shown in ball-and-stick representation and colored according to the chain to which they belong (blue and gold). Hydrogen bonds are shown as red-dashed lines. **(c)** Contacts between the N-linked glycan (with carbon atoms colored green, oxygen red, and nitrogen blue) and VSGsur, shown and colored as in (b). GlcNAc indicates N-Acetylglucosamine, MAN indicates mannose, and βMAN indicates β-mannose. **(d)** Schematic illustrating the contacts of VSGsur to suramin, focused on the ligand (generated by PDBsum^52^ using LigPlot^53^). Structures were illustrated with CCP4mg^48^.

Four amino acids from each monomer in the dimer contribute the bulk of protein contacts to the drug: H122, R125, H126, and R292 (Fig. 3, b and c). These positively charged residues are centered in the cavity of the dimer interface, burying the majority of surface area upon drug binding. These major contacts are buttressed with a set of minor interactions from residues L49, R119, A123, and A129, as well as from the N-linked glycan, several of these moieties only making contact from one protein monomer to some of the more distally positioned elements of suramin (such as the more poorly ordered sulfonic acids connected to the naphthalene rings, Fig. 3, c and d). The side chain nitrogen atoms from two arginine residues, R125 and R292, both make hydrogen-bonds with the sulfate oxygens of the most distal naphthalene moieties of suramin, as does H126. Other residues such as L49, R119, A123, and A129 primarily make close-approach, non-bonding contacts (Fig. 3d).

The interface centers on the imidazole rings of H122. In both monomers these residues make extensive contacts, stacking with two of the benzene rings of suramin (Fig. 3b). In the native protein structure, H122 has two different, partially occupied conformations in the monomer of VSGsur (Supplementary Fig. 7, a-c). One is very similar to the conformation seen in the suramin bound co-crystal structure whereas the other is a different rotamer, flipped away from this position. The presence of the drug locks the histidines in the stacking interaction with the suramin benzene rings, abolishing any evidence of the other possible conformation. The suramin benzene ring between the two rings contacted by H122, and which is centered in the dimer interface, interacts with VSGsur via backbone and sidechain contacts from R125 and H126. Interestingly, the benzene group of the compound used to phase the structure (5-amino-2,4,6-triiodoisophthalic acid, I3C or “Magic Triangle”, Methods), occupies the same position as the suramin benzene groups, stacking similarly with H122 (Supplementary Fig. 7d, Supplementary Table 1).

A secondary suramin binding mode was also discovered in a minority of the crystals. In this mode, H122 adopts the other possible conformer seen in the native structure. This conformation we term “closed” for how it reduces the size of the cavity (Supplementary Fig. 7b). In the closed conformation soak with suramin, the drug cannot occupy the same position as in the open conformation due to steric clash. Consistent with this, we observe difference density of a size possibly consistent with a molecule like suramin but positioned differently in a more extended conformation. However, the density is too weak to model with confidence. We hypothesize that as in the native structure, H122 is able to adopt two conformations, but that the open conformation with the stacking of the histidine rings over the suramin benzene groups leads to the better ordered interaction and thus likely represents the preferred binding mode. The closed conformation still leaves room for suramin to occupy the cavity, however, and may represent either a minor energetic state or transition from solution to the most stable binding mode.

The N-linked sugar groups (the N-acetylglucosamine (GlcNAc) rings, the β-mannose, and another mannose residue in the carbohydrate chain) make contacts to the sulfonic acids decorating the naphthalene rings (Fig. 3c). These contacts only occur for the sugar chain of one molecule, as the asymmetric positioning of suramin in the cavity leads the other naphthalene group to protrude too far from the carbohydrate to make effective contacts.

### Mutations in the suramin binding site abolish drug resistance

Altogether, the ITC and structural results show that unlike other VSGs, even the related VSG13, VSGsur binds suramin specifically and with high affinity. To examine whether the resistance phenotype of cells expressing VSGsur is tied to this binding, we created mutants (Supplementary Fig. 8, Methods) that should not disrupt the fold of VSGsur. One mutant was a direct loss-of-contact alteration, changing H122 to alanine (H122A), thereby removing the stacking of the two histidine rings to the benzene groups in the most ordered binding mode of the ligand. The second was indirect, mutating N130 to alanine (N130A), thereby preventing the N-linked glycosylation from occurring and removing the sugar-to-drug interactions.

Fig. 2a summarizes these results. Both mutants were viable and grew in the absence of the drug, although the N130A mutant evinced a lower growth rate with a doubling time of 8.9h compared to 6.5h for the wild type and 6.9h H122A mutant (Supplementary Fig. 4). The mutant VSGs behaved similarly to the wild type protein during purification, suggesting that the changes did not destabilize or otherwise compromise the proteins. However, they both lacked resistance to suramin (Fig. 2a). This loss is correlated with a loss in binding to the drug (Fig. 2b), directly linking the suramin-VSGsur interaction to the resistance phenotype.

To verify that the mutations did not perturb the fold of VSGsur, we solved the crystal structure of the VSGsur mutant H122A (Supplementary Table 3), showing a nearly identical structure to wild type VSGsur. We were unable to obtain well-diffracting crystals of the mutant N130A, likely as the sugar is significantly involved in crystal packing contacts. High concentration, long-duration soaks of suramin into the H122A mutant (Methods, Supplementary Table 3), however, revealed the presence of significant difference density in the pocket, but the density was poorly ordered and we were unable to confidently model suramin (Supplementary Fig. 9).

### Distal alterations to the suramin binding pocket are associated with increased resistance to suramin

We further subjected *T. b. rhodesiense* STIB900_sur^18^, which expresses VSGsur and is about 100-fold more resistant to suramin than other strains (IC_50_ of 1.1±0.13 µM), to increasing suramin pressure *in vitro* (up to 18 µM over one year). This produced “supermutants” that tolerated higher levels of the drug (IC_50_ of 11±1.2 µM). Analysis of the expressed VSG of a highly resistant clone showed that it was VSGsur with 14 mutations in the nucleotide sequence, producing eight amino acid substitutions in the protein sequence. These mutations clustered in the structure in two locations (Supplementary Fig. 10). G174T, E175A, and K288A are located around the N-linked glycan with several contacts to the sugar. They do not, however, contact the drug but could conceivably modulate contacts to suramin indirectly through its interactions with the carbohydrate. A304S, T313A, D317E, A318T, and A326T cluster on one face of the bottom lobe, distal from the drug binding site, creating a patch of surface of exposed residues. This distance from the binding pocket makes it very likely that these mutations do not affect drug binding and thus could represent a functional region of VSGsur that interacts with other factors critical to generating resistance.

A construct of the mutant VSGsur was targeted into the active VSG expression site of *T. b. rhodesiense* STIB900_sur1 (Supplementary Figure 11a). Of four transfected clones, two had replaced VSGsur with the mutant version and two had retained the original VSGsur, as verified by PCR. The “pseudo-clones” with the original VSGsur sequence showed equivalent resistance to previously characterized strains expressing VSGsur, whereas the two clones that expressed the mutant version of VSGsur showed an increase in their IC_50_ to suramin (Supplementary Fig. 11b), suggesting that the “supermutant” mutations indeed further enhance suramin resistance. However, this increased resistance was only 2-3-fold higher than wild type VSGsur, indicating that there are likely other mutations in the selected supermutant strains that synergize with the VSGsur mutations to produce the full 11-fold increase in IC_50_.

### Alterations in endocytic trafficking are not coupled to resistance

Wiedemar et al. had shown that VSGsur expressing strains have reduced levels of endocytic trafficking relative to VSG2 (also called VSG221), and, in particular, a reduced internalization of LDL, a factor hypothesized to be involved in suramin uptake into the cell^21–23^. Therefore, VSGsur might enhance resistance to the drug by decreasing suramin uptake through reducing endocytosis via the LDL receptor. However, as VSGsur likely comprises over 99% of the surface protein on the parasite and binds suramin with nanomolar affinity at blood pH, understanding suramin uptake in these strains is complicated by the potential for a massive influx of the drug as the VSG is internalized and then recycled back to the membrane^31^. To interrogate this model, we performed endocytosis experiments with blue dextran and bodipy-labeled LDL (Fig. 4, Methods). Our data show that, consistent with previous results^23^, there is no generalized defect in trafficking in our VSGsur expressing strains (Fig. 4, a and b). In fact there appears to be a wide variation between different VSGs on the rates of trafficking (Fig. 4a). While we do observe that VSGsur cells have a significant decrease in LDL uptake compared to VSG2 (although, again, there is substantial variance among different VSGs, Supplementary Fig. 10), loss-of-binding suramin mutants show no significant alteration in these kinetics (Fig. 4c). Similar results are seen with uptake via the transferrin receptor (Fig. 4d). As these mutants also lose resistance to suramin, it seems that the dynamics of LDL uptake (and thus, presumably, suramin via this pathway) are not strongly coupled to the resistance phenotype in VSGsur. In contrast, the key determinant appears to be the binding of the drug to the VSG.

**Fig. 4:**
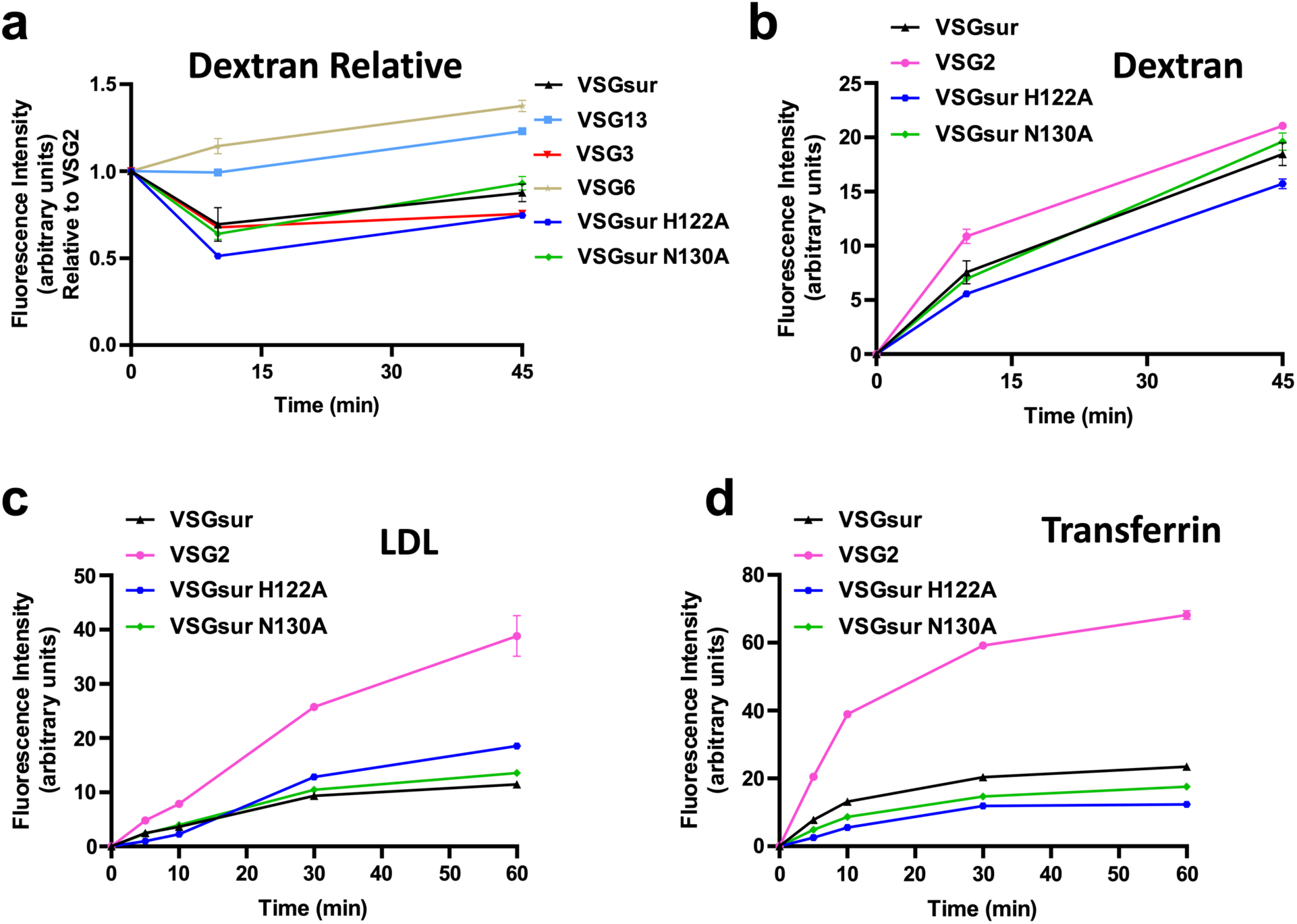
VSGsur and the loss of contact mutants have indistinguishable endocytic kinetics. **(a)** Alexa 488-dextran endocytosis by *T. b. brucei* 2T1 cells expressing a variety of different VSG genes. Due to technical limitations, each cell line could not be analyzed simultaneously within each experiment for each ligand. Therefore, VSG2 was used as a control in all experiments, allowing the calculation of the relative uptake rate for each cell line rate as compared to the uptake rate of each ligand observed by VSG2 expressing cells within each separate experiment. Each of these graphs therefore represent the combined experimental results from 2 to 3 separate experiments. Cell lines with a relative fluorescence intensity below 1 at a given time point have less efficient endocytic rates compared to VSG2 expressing cells, and vice versa. All graphs share the same Y axis. Error bars when present denote the range (N=2 experimental replicates for each cell line at every time point), while error bars were left out when they were smaller than the size of the data point and therefore not visible. **(b)-(d)** Alexa 488-dextran (b), bodipy-LDL (c), and Alexa 488-transferrin (d) endocytosis by *T. b. brucei* 2T1 cells expressing VSGsur (black triangles), VSG2 (pink circles), VSGsur H122A (blue hexagons), or VSGsur N130A (green diamonds). All graphs depict arbitrary fluorescence units, determined by flow cytometry, on the Y axis. Each graph represents one of multiple biological replicates of each experiment, with N=2 experimental replicates for each time point. Error bars when present denote the range, while error bars were left out when they were smaller than the size of the data point and therefore not visible.

## Discussion

Despite the wide genetic variance in the VSGs, only a handful of the membrane distal, large NTD domains have been characterized at the atomic level^24–27^. The recent crystal structure of VSG3 revealed topological divergence from the folds of other VSG NTDs, demonstrating that there is significantly more variation in architecture of this protein family than previously considered^25^. This structure also uncovered an unexpected post-translational modification (*O*-linked sugar at the top of VSG3) that strongly modulates the host immune response. With the structures of VSGsur and VSG13 presented here, we show that the “antigenic space” of the VSGs is much broader than anticipated even from the VSG3 results. Moreover, the co-crystal structure of VSGsur with the trypanocidal compound suramin directly links the binding of the drug to the resistance phenotype displayed by strains of *T. brucei* expressing VSGsur.

This binding of suramin establishes that the VSGs can have a function beyond that of antigenic variation. This idea is buttressed by recent results that show that VSG2 is able to bind a ligand as well (the divalent metal calcium, manuscript in preparation). Also consistent with this, the recently determined crystal structure of the *T. congolense* haptoglobin–hemoglobin receptor^30^ shows that this protein adopts a fold very similar and clearly related to the VSGs, packing functionally with all possible VSGs on the surface as it is required for parasite survival in the host to scavenge iron. These observations raise the possibility of other functions for the VSGs in binding small molecules or even proteins, and to therefore function for the benefit of the pathogen in manners more diverse than mere immune evasion.

We are not aware of any data showing that suramin-resistant field isolates express VSGsur. VSGsur was identified in the laboratory under cell culture conditions of increasing suramin concentrations, where there is no pressure to switch to alternative VSGs that can prevent immune clearance (in contrast to the situation in the host). If VSGsur undergirds resistance in the context of infection, antigenic variation and VSG switching would need to be modeled. Wiedemar et al.^18^ noted that short-term survival under low-dose suramin treatment in animals could allow for the development of alternative mechanisms of drug resistance. Another model would involve the display of multiple, antigenically distinct versions of VSGsur. This could conceivably occur through gene duplication and divergence of surface features while preserving the suramin binding cavity or through the formation of mosaics that switch CTDs in a manner that alters the antigenicity of the NTDs (as has been recently reported^32^).

While the binding of suramin to VSGsur is seen to be required for drug resistance, it is still unclear by what mechanism VSGsur is producing this effect. Genetic studies coupled with examination of the trypanosome’s single lysosome suggest that suramin accumulates in this compartment, while other evidence links suramin toxicity to effects on the glycosome and impairment of cytokinesis^19,20^. Our internalization data decouple suramin uptake via the LDL endocytic pathway from the VSGsur-related mechanism of resistance. This focuses attention on the interaction of the drug with VSGsur itself. However, it is unclear how to link resistance to (presumably) lysosomal trafficking and downstream toxicity to the VSGsur-suramin interaction. The simplest model involves VSGsur acting as a cellular repository or “sponge” that reduces the trafficking of the drug, shunting it to some kind of disposal pathway and thereby reducing the effective concentration. Since VSGsur will be produced at high levels in the trypanosome and exported toward the cell surface from the Golgi through the vesicular trafficking pathways, it is conceivable that suramin that is internalized through other pathways (e.g., via the LDL receptor and ISG75) could comingle with VSGsur in specific endocytic compartments, the latter binding the drug and preventing it from trafficking further (Fig. 5). Such a disposal pathway could involve shuttling the drug from the vesicular trafficking pathway to the extracellular space or another compartment that renders it inactive.

**Fig. 5:**
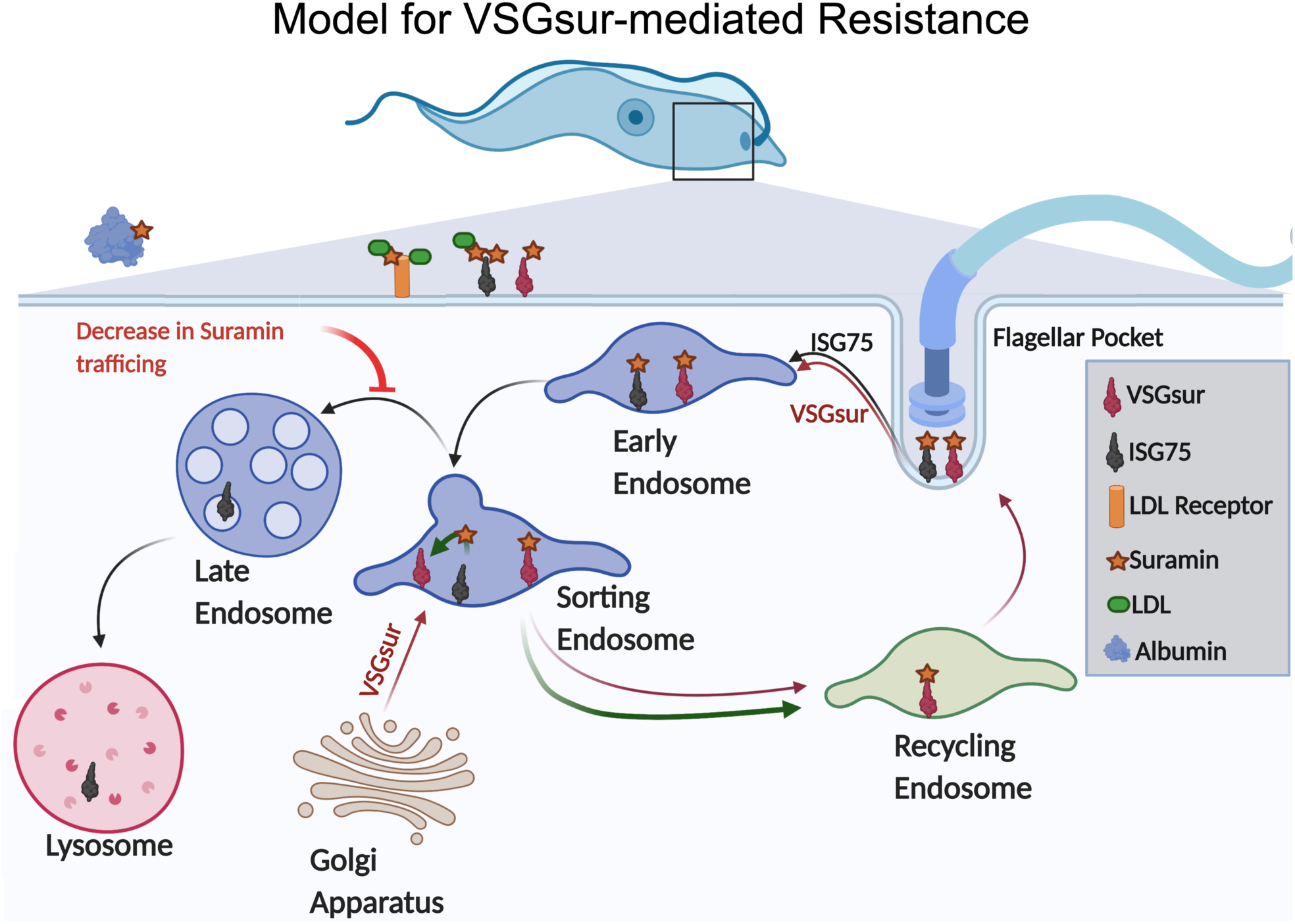
Model for possible mechanism of VSGsur-mediated resistance to suramin. A schematic illustration of how VSGsur could shuttle suramin away from toxic endocytic internalization pathways. A single trypanosome cell is depicted in blue at the top of the diagram, the boxed region magnified showing the vesicular transport compartments. Critical players like ISG75, the LDL receptor, blood proteins, and VSGsur are shown as defined in the key. In this model, VSGsur acts as a “sponge” to remove suramin from the endocytic pathway, decreasing its effective concentration and thereby rendering the cell more resistant to the drug.

The VSGsur “supermutants” could offer a tantalizing clue toward solving this puzzle and uncovering the mechanism. While one subset of the mutations clusters around the carbohydrate and could conceivably alter binding to the drug, another set clusters in a surface patch on the bottom lobe of the protein. This patch is suggestive of a binding site and may point the way toward the macromolecular interactions involved in suramin resistance. Future experiments will likely require a tracking of the drug through the trypanosome to establish how VSGsur binding modulates toxicity. The results of this study focus such investigations squarely on the VSG as the key player in resistance, and also provide numerous tools (such as structure based mutants) to interrogate the biochemical process allowing strains to survive in high concentrations of suramin.

## Methods

### Cloning and Production of *T. brucei* strains

A VSG13 expressing *T. brucei* clone was produced as described^33^ and was kindly provided by the group of Luisa Figueiredo (Universidade de Lisboa). VSGSur from *T. brucei rhodesiense* (GenBank: MF093647.1) was codon-optimized and synthesized as a pUC19 clone (BioCat, Germany), and introduced into *T. brucei* strain Lister427 as described in Supplementary Methods. VSGsur mutants N130A and H122A were generated by site-directed mutagenesis using the QuikChange Lightening kit (Agilent Technologies) according to the manufacturer’s protocol. Transfections were performed into a *T. brucei brucei* cell line expressing VSG2, termed 2T1^34^. 5-10 micrograms of purified plasmids were mixed with 100 µl 3 ×10^7^ cells in Tb-BSF buffer^35^ (90 mM Phosphate buffer, pH 7.3, 5 mM KCl, 50 mM HEPES, pH 7.3, 0.15 mM CaCl2) and electroporated using a Lonza Nucleofector 2b, program Z-001 or X-001. After 8 hours of incubation hygromycin B was added to a concentration of 5-25 µg ml^-1^. Single cell clones were obtained by serial dilutions in 24 well plates in standard culture medium and screened by FACS for VSG2-antibody binding. VSG2-antibody negative clones (positive transfectants were identified by lack of binding) were sequenced: RNA was isolated using the RNeasy Mini Kit including an on-column DNase treatment (Qiagen). Complementary DNA was synthesized with SuperScript IV Reverse Transcriptase (Invitrogen) according to the manufacturer’s protocol. Amplification was performed with a forward primer binding the spliced leader sequence and a reverse primer binding in the VSG 3’UTR using Phusion high fidelity DNA polymerase. PCR products were purified by gel extraction from a 1% agarose gel, followed by the NucleoSpin Gel and PCR clean up kit (Macherey-Nagel) and sent to Eurofins (Ebersberg) for sequencing using the same primers as for the PCR.

### Purification of VSGs

*T. b. brucei* expressing VSG2, VSG3, VSG13, VSGsur, VSGsur-N130A, and VSGsur-H122A mutants were cultivated *in vitro* in HMI-9 media (formulated as described by PAN Biotech without FBS, L-cysteine, or ß-mercaptoethanol), supplemented with 10% fetal calf serum (Gibco), L-cysteine, and ß-mercaptoethanol. Cells were cultured at 37°C and 5% CO_2_. VSGs were purified according to established protocols^37^. Briefly, cells were pelleted and then lysed in 0.4 mM ZnCl_2_. The lysis mixture was centrifuged and the pellet containing the membrane material was resuspended in pre-warmed (40 °C) 10 mM phosphate buffer, pH 8. Following a second centrifugation, supernatant containing VSG protein was loaded onto an anion-exchange column (Q-Sepharose Fast-Flow, GE Healthcare), which had been equilibrated with 10 mM phosphate buffer, pH 8. The flow-through containing highly pure VSG was concentrated in an Amicon Stirred Cell and optionally shock frozen in liquid nitrogen and stored at −80°C.

For crystallization of VSG13, the C-terminal domain was removed by digestion with Endoproteinase LysC (NEB) at a 1:800 LysC:substrate ratio by mass for 1-2h at 37°C. VSG13 N-terminal domain was purified by gel filtration on a HiLoad 16/600 Superdex 200pg column (GE Healthcare) in 50 mM HEPES, pH 7.6, 150 mM NaCl. Lysine residues were methylated (as per^38^), using Dimethyl-amine-borane complex (Sigma) and Formaldehyde (Thermo, Methanol-free). The methylated VSG13 N-terminal domain was purified by gel filtration on a Superdex 200 Increase 10/300 GL (GE Healthcare) in 10 mM Tris.Cl, pH 8.0. Fractions from the gel filtration runs were subjected to SDS–PAGE analysis for visual inspection.

For crystallization of VSGsur and VSGsur mutants, the C-terminal domain was removed by limited proteolysis with trypsin at a 1:100 trypsin:substrate ratio by mass for 1h on ice. The VSGsur N-terminal domain was purified by gel filtration on a HiLoad 16/600 Superdex 200pg column (GE Healthcare) in 10 mM Tris.Cl, pH 8.0. The peak fractions were pooled and concentrated to 4 mg/ml.

### Structural Determination of VSG13

Purified methylated VSG13 N-terminal domain was concentrated to 2.5 mg ml^−1^ in 10 mM Tris.Cl pH 8.0. Crystals were grown at 23°C by vapour diffusion using hanging drops formed from mixing a 1:1 volume ratio of the protein with an equilibration buffer consisting of 1.8-2.0 M (NH_4_)_2_SO_4_, 100 mM Tris.Cl pH 8.5. For cryoprotection, crystals were transferred into the same buffer including 20% glycerol and flash-cooled immediately afterward to 100 K (−173.15 °C). For phase determination, crystals were soaked for 30s in a buffer containing 0.5 M sodium bromine and 20% glycerol before flash-freezing.

Native data were collected at the European Synchrotron Radiation Facility (ESRF) at a wavelength of 1Å on beamline ID29 and bromine soaks at the Diamond Light Source at 0.9198Å on beamline i03 (Supplementary Table 2). The data were phased by single-wavelength anomalous diffraction (SAD) from 33 bromine sites that were identified using the SHELX suite^39^. Several models were manually combined from automated model building by PHENIX^40^ and CRANK of CCP4^41^ into maps produced from the SHELX sites and improved by several cycles of manual building, auto-building (PHENIX), and refinement (PHENIX) into the native dataset (Supplementary Table 2).

### Structural Determination of VSGsur and VSGsur mutants

Crystals were grown at 4°C by vapour diffusion using hanging drops by mixing 2 µl of the protein solution (4mg/ml) with 2 µl of the equilibration buffer (19-24 % PEG 400, 100 mM TEA/HCl pH=7.5 and 10 % (v/v) isopropanol). The crystals were transferred to a cryobuffer (40% PEG400 and 100 mM TEA/HCl) and flash-cooled in liquid nitrogen. Native datasets were collected at a wavelength of 0.9184 Å at the Helmholtz-Zentrum Berlin at Beamline MX 14.1. For phasing the crystals were soaked overnight in 50 mM 5-amino-2,4,6-triiodoisophthalic acid (I3C)/LiOH (Magic Triangle, Jena Biosciences)^42^ in cryobuffer. The I3C soaked crystals were collected at a wavelength of 2.066 Å at the Helmholtz-Zentrum Berlin at Beamline MX 14.2. The structure was solved by SAD from the anomalous signal from two I3C molecules bound per VSGsur monomer using the SHELX^39^ and the HKL3000^43^ suite. Arp/wARP^44^ was used for automated building of the initial model. PHENIX^40^ and COOT^45^ were used for several cycles of model building and refinement.

Suramin complexes were obtained by either soaking VSGsur crystals in the drug or by pre-mixing and growing crystals *de novo*. Complexes were produced by soaking the crystals between 1 hour and 6 days in cryobuffer supplemented with 0.77 - 7.7 mM Suramin before cryo-cooling. Co-crystals were obtained at 4°C using 3 mg/ml VSGsur, 0.7 mM Suramin and an equilibration buffer containing 16-20% PEG 3350, 200 mM NaCL and 100 mM Hepes/NaOH pH=7.5. Prior to cryo-cooling these crystals were transferred to 16 % PEG 3350, 200 mM NaCl, 100 mM Hepes/NaOH pH=7.5, 0.7 mM Suramin and 25 % PEG 400. The VSGsur-suramin datasets were collected at the Helmhotz-Zentrum Berlin, the Paul Scherrer Institut Villingen, and Diamond Light Source. The structures were solved by molecular replacement with PHENIX using the native structure of VSGsur as the search model (Supplementary Table 1).

The incubation of native crystals with suramin produced an unexpected effect on the crystal packing. Native VSGsur crystallized in the space group P2_1_2_1_2 with a single molecule in the asymmetric unit and a two-fold axis of crystallographic symmetry aligned with the dimerization axis, thereby producing a crystallographic dimer highly similar to that observed in other VSGs. One hour soaks of native crystals in 0.7 mM and 7.7 mM suramin did not change these parameters and led to structures where additional difference density appeared in the cavity but which could not be modeled. However, as the soak time increased (greater than 4 hours), a shift in the crystallographic symmetry was observed and correlated with increased evidence of electron density for suramin. In particular, the two-fold rotational symmetry no longer aligned with the dimerization axis of rotation, causing a shift in the contents of the crystal to harbor two molecules (a single homodimer of VSGsur) in the asymmetric unit in a lower symmetry space group. These changes were tolerated by the crystals, maintaining high diffraction. In these soaks, electron density for suramin was clear and able to be modeled.

### Uptake Assays

The LDL and transferrin uptake assays were performed as described^23^ with slight modifications while the dextran assays were performed similarly. Cells were cultured to a density of approximately 1 ×10^6^ cells per mL of HMI-9. For LDL and transferrin, the cells were pelleted and washed in serum free medium containing 1% BSA before being returned to 37°Cfor 15 minutes to remove any surface-bound ligand. Approximately 100,000 cells were aliquoted into assay tubes in duplicate and kept at 37°C. Pre-warmed bodipy-labeled LDL (Invitrogen; ThermoFisher catalog number I34359) suspended in serum free medium containing 1% BSA was added to the tubes at the indicated time points such that the final volume of the assay was 150 µL and the concentration of LDL was 15 µg/mL. Being at a much higher concentration as a stock, 2 µL of transferrin (Invitrogen; ThermoFisher catalog number T13342) was added to the assay tubes to a final volume of 200 µL and a final concentration of 50 µg/mL. The assays were conducted such that all conditions reached the end point of the assay simultaneously. At time zero, all vials were placed on ice and 1 mL of ice cold serum free medium containing 1% BSA was added to the tubes. The cells were washed twice by centrifugation at 2000 x G for 5 minutes and resuspended in ice cold serum free medium containing 1% BSA. The vials were then kept on ice until being analyzed on a Becton Dickinson FACSCalibur. Fluorescence intensity was quantified by analysis in FlowJo after gating on cells determined by forward and side scatter. The geometric mean of fluorescence was then calculated for between 7000 and 10,000 cells per replicate per time point for each cell line assayed. To control for differences in intrinsic auto-fluorescence, each cell line was individually normalized to its own “time 0” point, which was collected from cells that had been exposed to ice cold ligand and immediately placed on ice. These “time 0” points were generally within only 1-2 fluorescence units of each other. For dextran, approximately 100,000 cells were aliquoted into assay tubes and pelleted. The cells were suspended in 50 µL of 0.5 mg/mL Alexa 488-dextran diluted in HMI-9 and kept at 37°C until reaching the expiry of the incubation. Once reaching those time points, the cells were immediately placed on ice and 1 mL of ice cold HMI-9 was added to the tubes. The cells were washed three times by centrifugation at 2000 x G for 5 minutes and resuspended in ice cold HMI-9. The vials were then kept on ice until being analyzed on a Becton Dickinson FACSCalibur and fluorescence was quantified as above.

### Production of “Supermutants”

Bloodstream-forms of *T. b. rhodesiense*_sur1 were further selected *in vitro* by stepwise increases of the suramin concentration from 1 µM to 18 µM over the course of one year. The expressed VSG gene was amplified from cDNA by PCR with primers binding to the 5’ spliced leader and a conserved sequence (gatatattttaaca) on the 3’ untranslated region of VSGs^46,47^. For transfection, a construct was used containing the blasticidine resistance gene and the coding sequence of mutant VSGsur separated by a αβ tubulin splice site, and framed with the 5’ and 3’ UTRs of VSGsur (Supplementary Fig. 11). Transfectant pseudoclones were generated by limiting dilution as described^18^.

### Determination of growth rates and drug sensitivity assays

Cumulative growth curves for different VSG-expressing *T. brucei* strains were determined by dilution of the cells to 10^5^ ml^-1^ in 25 cm^2^ cell culture flasks in duplicate. After incubation with 0.7µM suramin for 24 h the cell densities were determined by cell counting using a Neubauer hemocytometer. This procedure was repeated for 3-4 days in a row. Analysis was performed with GraphPad Prism, using a nonlinear regression model for curve fitting (Exponential growth with log).

For the drug sensitivity assays, serial dilutions of suramin (Sigma, dissolved in water and stored in aliquots at −20°C) as well as blank cultures without suramin, were prepared in 24 well plates (at least 3 technical replicates). Cells were added to a density of 10^5^ ml^-1^. After 24 h the cell densities were determined with a Neubauer hemocytometer. The 50% Inhibitory concentrations (IC_50_) were calculated with GraphPad Prism, using the non-linear regression model to fit the dose-response curve (variable slope; four parameters).

### Isothermal Titration Calorimetry (ITC)

ITC experiments were performed using a PEAQ ITC (Malvern) at 20°C. Titration buffers contained 10mM NaPi (pH 8.00) and 150mM NaCl. Proteins were transferred into the titration buffer by gel filtration chromatography, followed by concentration in 10k disposable ultrafiltration centrifugal devices. Protein concentrations were measured by UV absorbance at 280 nm. In each experiment, the protein concentration in the cell varied between 40-55µM. Suramin was injected at concentrations between 300 and 600 µM, and all samples were degassed prior to each experiment. All VSG protein-suramin experiments were performed at least in duplicate to check the reproducibility of the data. The data were baseline corrected, integrated, and analyzed with the PEAQ ITC Analysis software (Malvern), fitted using a single-site binding model.

### Data Availability

Coordinates and structure factors have been uploaded to the RCSB PDB (www.rcsb.org): VSGsur I3C (PDB ID 6Z79), VSGsur WT native (PDB ID 6Z7A), VSGsur + Suramin (PDB ID 6Z7B), VSGsur H122A (PDB ID 6Z7C), VSGsur H122A 0.77mM Suramin (PDB ID 6Z7D), VSGsur H122A 7.7 mM (PDB ID 6Z7E), VSG13 NaBr (PDB ID 6Z8G, and VSG13 native (PDB ID 6Z8H).

## SUPPLEMENTARY FIG.S AND TABLES

Figures 1-12

Table 1-3

## ACKNOWLEDGEMENTS

We acknowledge time at the European Synchrotron Radiation Facility (ESRF, beamline ID29, proposal MX1975, Gianluca Santoni and colleagues), the Diamond Light Source (DLS, beamline i03, proposal number mx18989-1, Neil Paterson and colleagues), the Helmholtz Zentrum Berlin (BESSY, beamline 14.1 and 14.2, proposal number MX-191-00036 and MX-192-00114, staff member Manfred Weiss and colleagues, specially Christian Feiler for the help with processing and solving the I3C SAD dataset), and the Paul Scherrer Institut, Villingen, Switzerland (SLS, beamline X06DA, proposal number 20182345, 20191097 and 20191895, Vincent Olieric and colleagues). We thank Luisa Figueiredo for supplying us with the VSG13 expressing *T. brucei* strain.

## AUTHOR INFORMATION

H.H. cloned VSGsur and M.vS. produced VSGsur mutants. S.H., N.W., and P.M. produced VSGsur “supermutant” clones and assays. M.vS. and J.Z. purified all VSGs. M.sV. crystallized VSG13. M.vS. and J.Z. crystallized wild type and mutant VSGsur proteins. C.E.S. phased and produced the first model of VSG13 and A.H. built and refined the structures of VSG13 native and the sodium bromide soak. J.Z. solved all crystal structures of VSGsur. P.J. worked with J.Z. on the refinement of VSGsur bound to suramin. M.vS. performed the suramin growth and toxicity assays. K.P. conducted ITC experiments. J.V. conducted endocytosis assays.

