## Supplementary Figures, Tables, and Methods for "A Parasite Coat Protein Binds Suramin to Confer Drug Resistance"

[1]Division of Structural Biology of Infection and Immunity, German Cancer Research Center [2]Division of Immune Diversity, German Cancer Research Center, Heidelberg, Germany. [3]Protein Expression and Purification Core Facility, EMBL Heidelberg, Meyerhofstraße 1, Heidelberg, Germany [4]Department of Molecular Biology, Princeton University, Princeton, New Jersey, USA [6]University of Basel, Basel CH-4001, Switzerland [6]Swiss Tropical and Public Health Institute, Basel CH-4002, Switzerland.

[*]These authors contributed equally to this work.

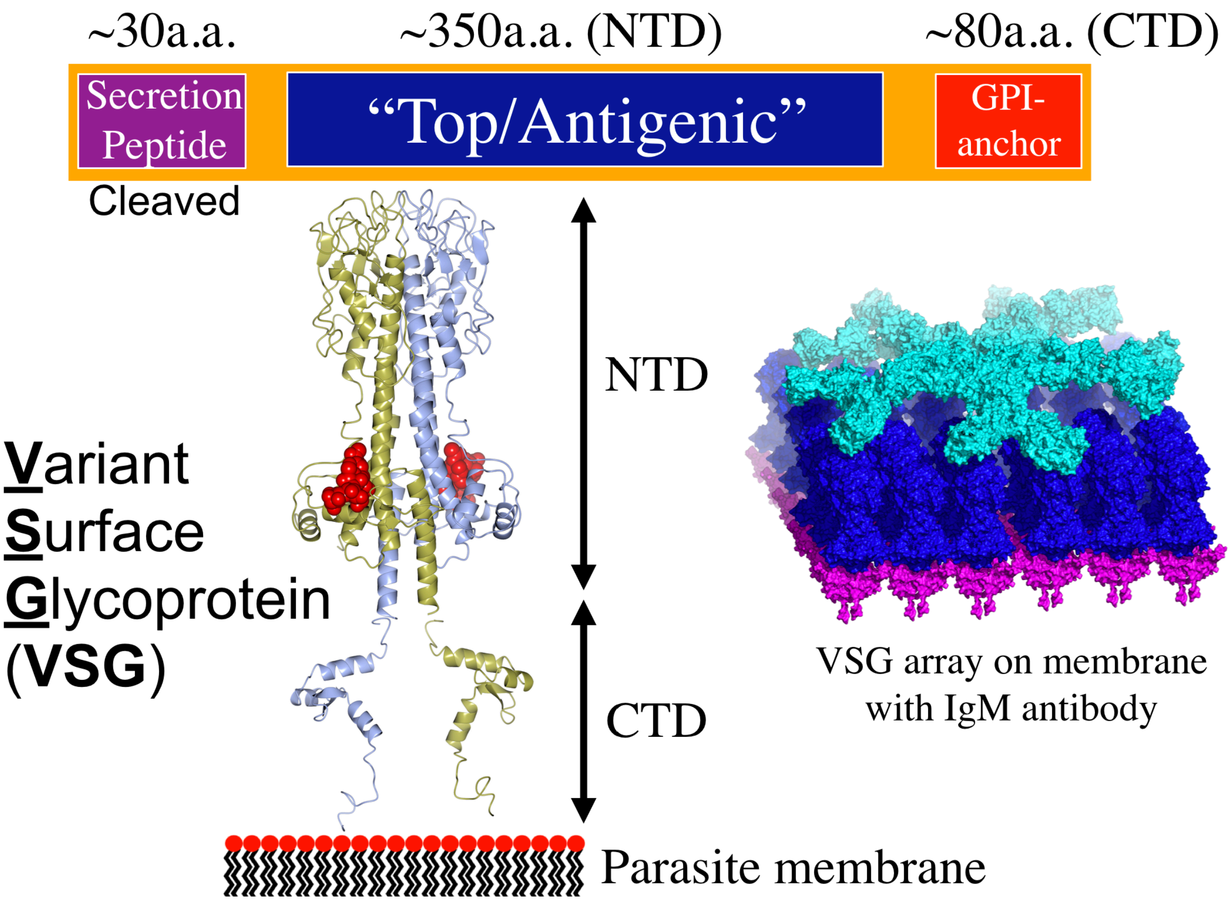


**Supplementary Fig. 1: Domain structure of the VSG proteins.** The top bar shows a schematic of the domain structure of the VSG coat proteins with rough amino acid size associated with each. NTD is the N-terminal domain and CTD is the C-terminal domain. Underneath is the structure of VSG1 schematically represented by a composite model of N and C-terminal domain structures (PDB IDs 5LY9 and 5M4T, respectively), illustrated over a cartoon of the parasite membrane. On the right is a model of an array of VSG proteins (NTD in blue, CTD in magenta) on the surface and interacting with a hypothetical model of immunoglobulin M (IgM shown in cyan).

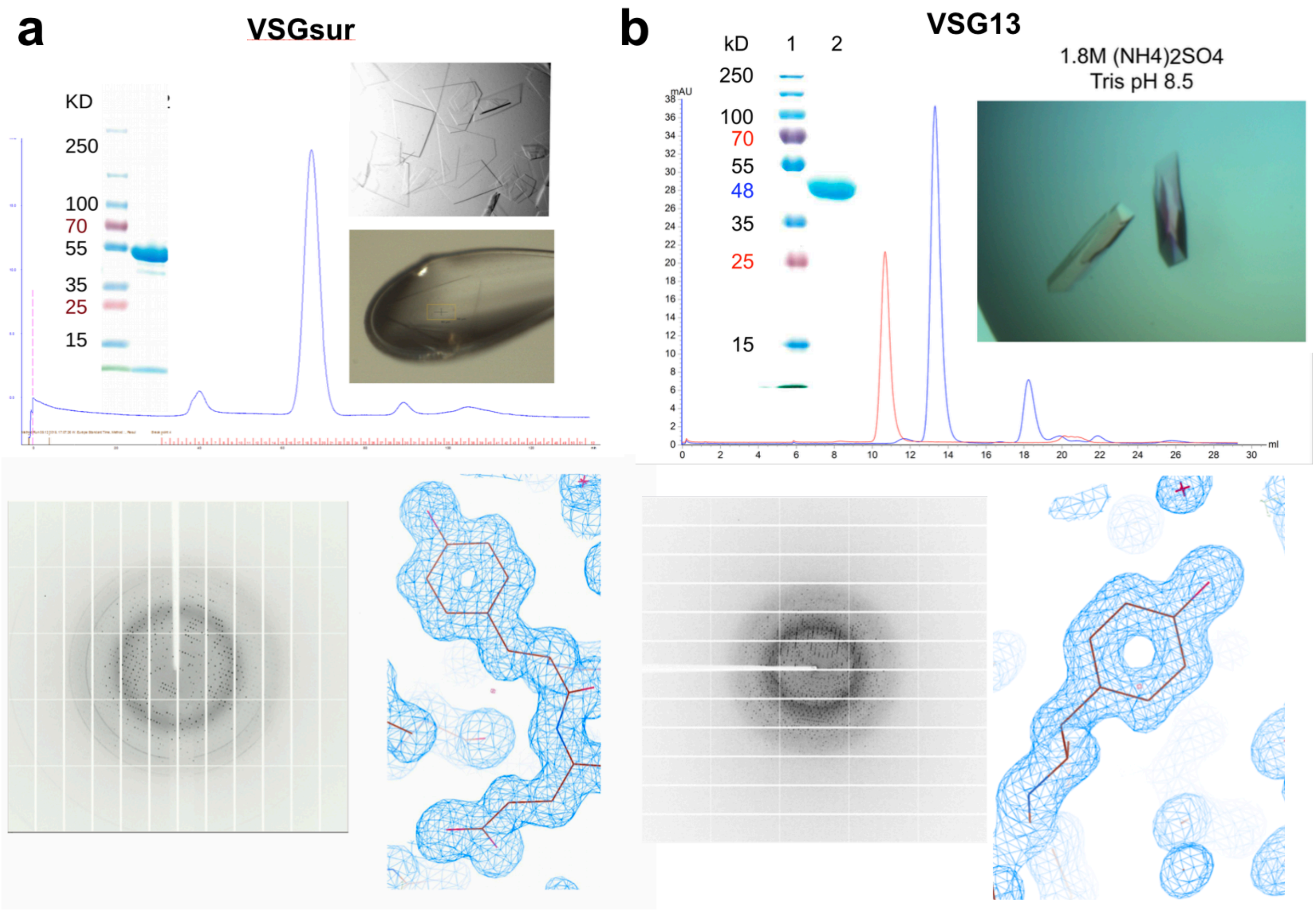


**Supplementary Fig. 2: Purification, crystallization, and representative electron density of VSGsur and VSG13.** Summary of various steps in the crystallographic structural solution of VSG13 and VSGsur. **(a)** Panels showing the gel filtration chromatogram (Superdex 200, Methods) of purified VSGsur NTD and a coomassie stained SDS-PAGE gel of the final material used for crystallization. Images of cryo-crystals grown in hang-drops and cooled crystals in the X-ray beam, a diffraction image, and final model 2Fo-Fc electron density contoured at 1σ. **(b)** Panels included are: the gel filtration chromatogram (Superdex 200, Methods) of purified VSG13 NTD (native in blue and reductively methylated in red, the latter running larger as perhaps a tetramer but occurring as a dimer in the asymmetric unit of the crystal), a coomassie stained SDS-PAGE gel of the final material used for crystallization, images of crystals prior to harvesting in hanging drops, a diffraction image, and final model 2Fo-Fc electron density contoured at 1σ.

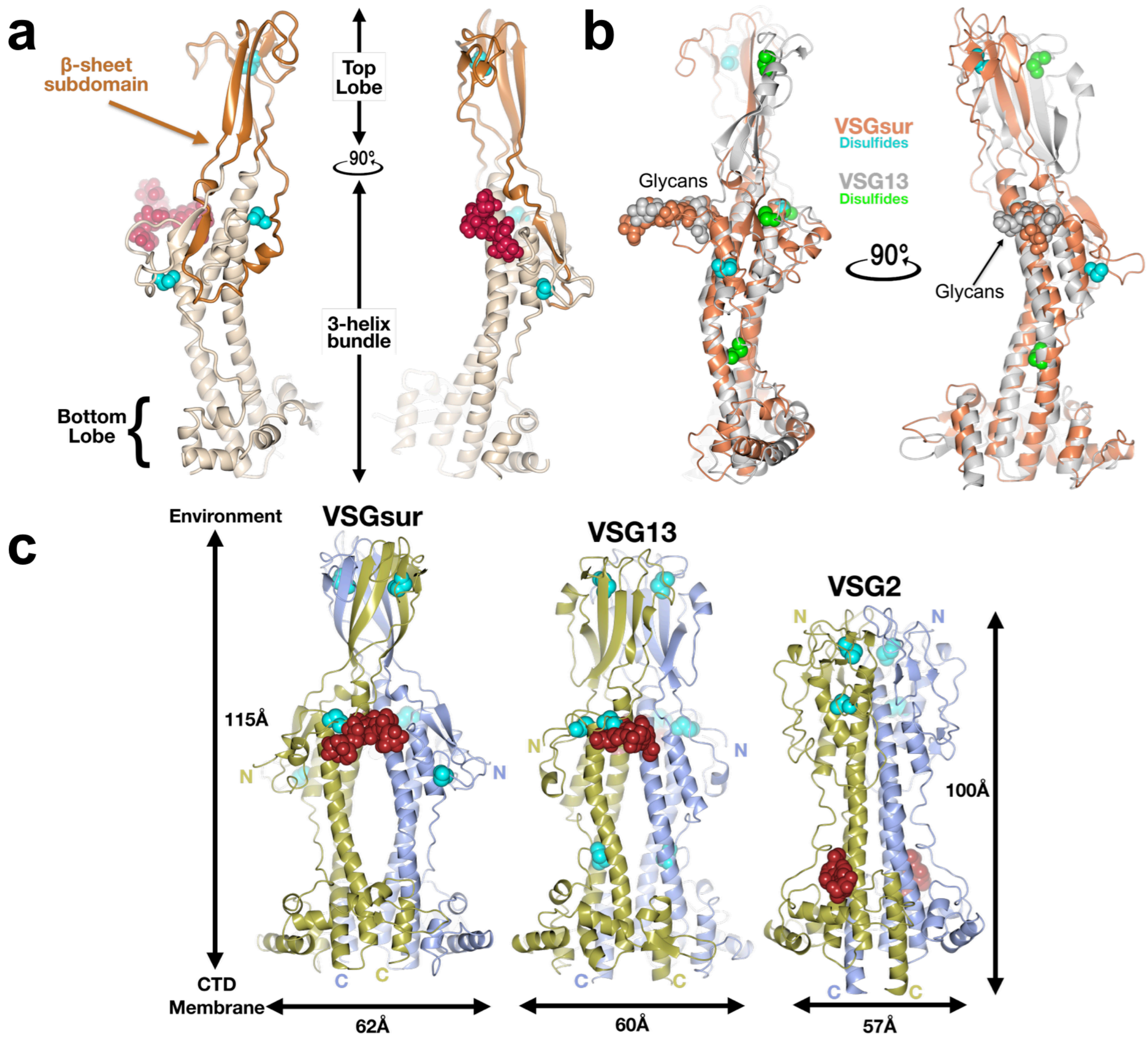


**Supplementary Fig. 3: Comparison of the VSG NTDs**

**(a)** A single monomer of the VSGsur NTD is shown as a ribbon diagram. The β-sheet subdomain that forms the top lobe is colored in orange and the rest of the structure in beige. The N-linked glycan is shown as red space-filling spheres, the cysteine disulfide atoms as space-filling cyan spheres. **(b)** Structural alignment of monomers of VSGsur (orange) and VSG13 (gray) with corresponding glycans and disulfides shown in space filling representation and colored the same as the protein to which they are linked. The alignment produces an overall root-mean-square deviation of 1Å for the conserved portions of the structure (calculated over 220 Cα positions that primarily encompass the three-helix bundle and elements of the bottom lobe). **(c)** Comparison of the structures of VSGsur, VSG13, and VSG2. The N-linked glycans are displayed as red space-filling atoms and the disulfide bonds are shown in cyan. Approximate dimensions of the molecules are noted, as well as the directions toward the external environment and toward the C-terminal domain (CTD) and plasma membrane of the trypanosome.

WT and Mutant VSGsur growth curves

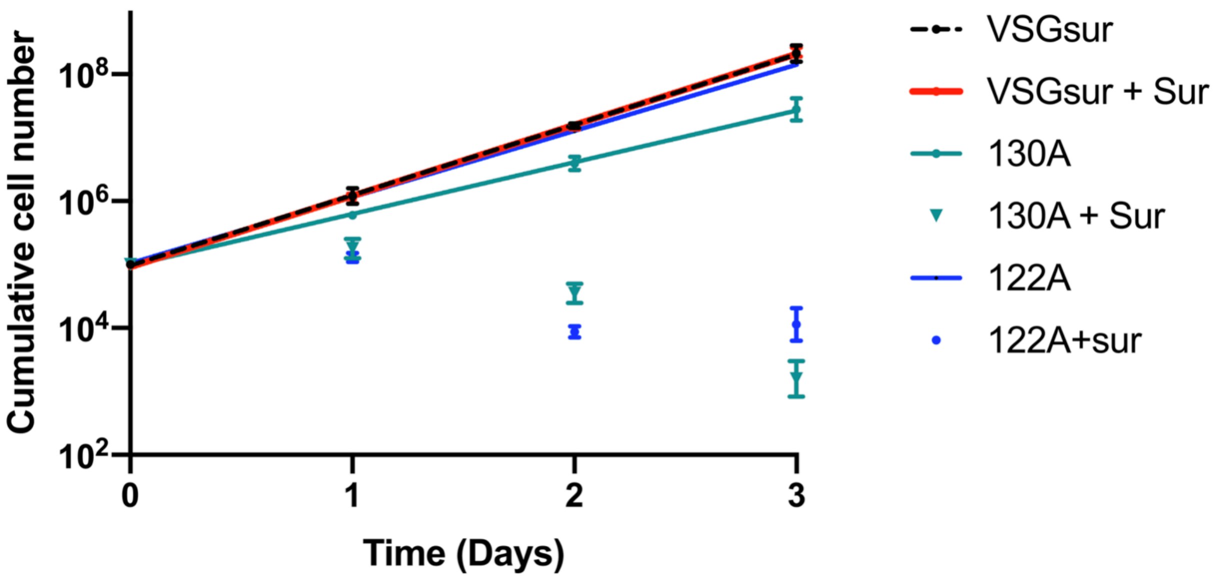


**Supplementary Fig. 4: Growth curves of VSGsur and VSG mutants.**

Strains expressing VSGsur and VSGsur mutants were grown with and without suramin (incubation of 0.7µM suramin for 24 h, Methods). The cell densities were determined by cell counting using a Neubauer hemocytometer. This procedure was repeated for 3-4 days in a row. Analysis was performed with GraphPad Prism, using a nonlinear regression model for curve fitting (Exponential growth with log).

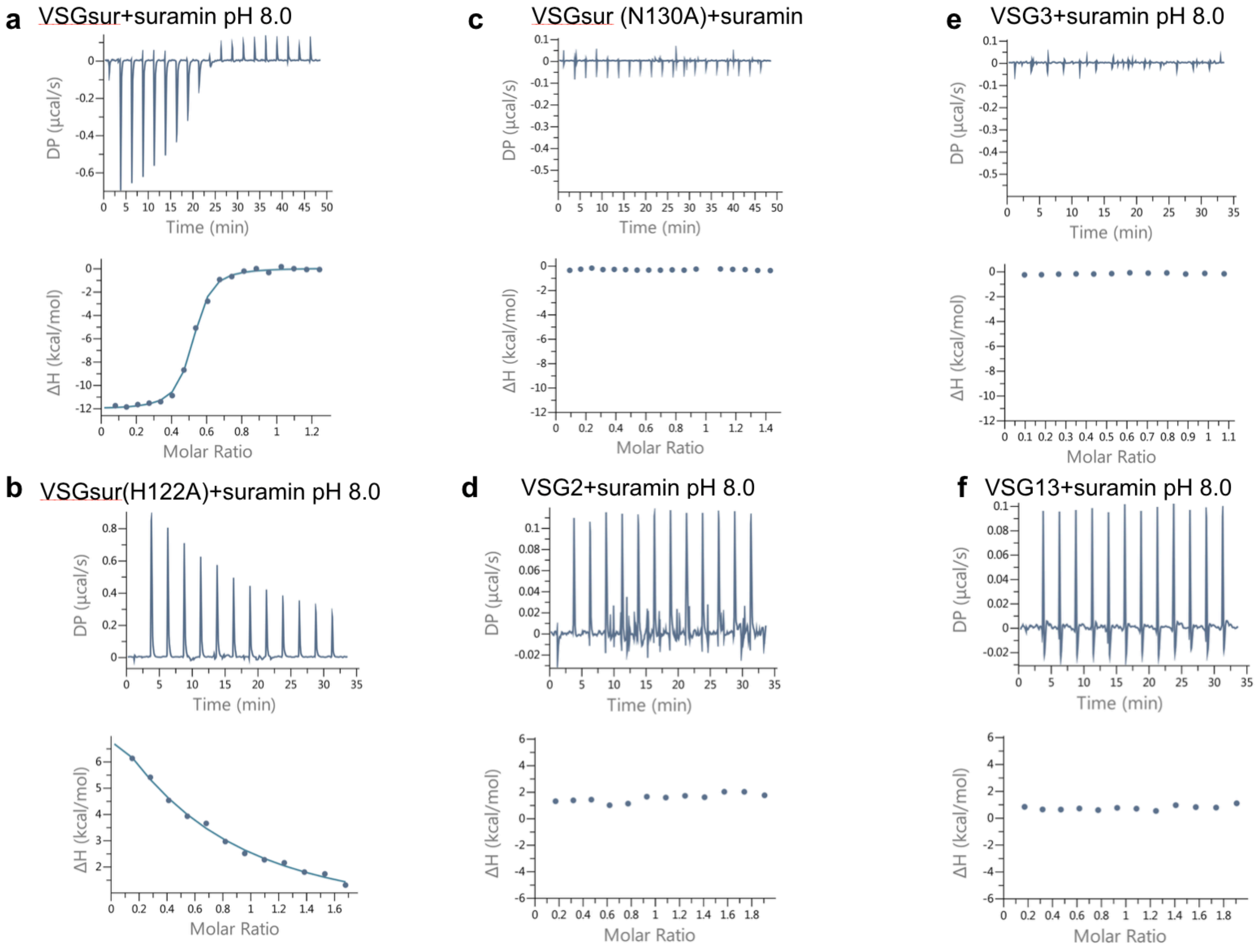


**Supplementary Fig. 5: Sample ITC results.**

ITC data for suramin binding to each VSG protein. The upper panels contain the baseline corrected raw data, and the lower panel contains the peak-integrated, concentration normalized data for the heat of reaction vs. molar ratio of suramin per VSG protein. **(a)** VSGSur, pH 8.0: 300µM suramin was titrated into 46µM VSG Sur pH 8., fitted with a single binding site model to calculate a Kd of 234+/- 28 nM and N of 0.49 +/- 0.03 **(b)** VSGSur H122A: 450µM suramin was titrated into 51.1µM VSGSur H122A. A Kd was not fit to the data, although it is clear that the mutation negatively affected the binding affinity. **(c)** VSGSur N130A: 300µM suramin was titrated into 40µM VSGSur N130A. No binding was detected. **(d)** VSG2: 200µM Suramin was titrated into 20µM VSG2 protein. No binding was detected. **(e)** VSG3: 300µM suramin was titrated into 53.1µM VSG3. No binding was detected. **(f)** VSG13: 200µM Suramin was titrated into 20µM VSG132 protein. No binding was detected.

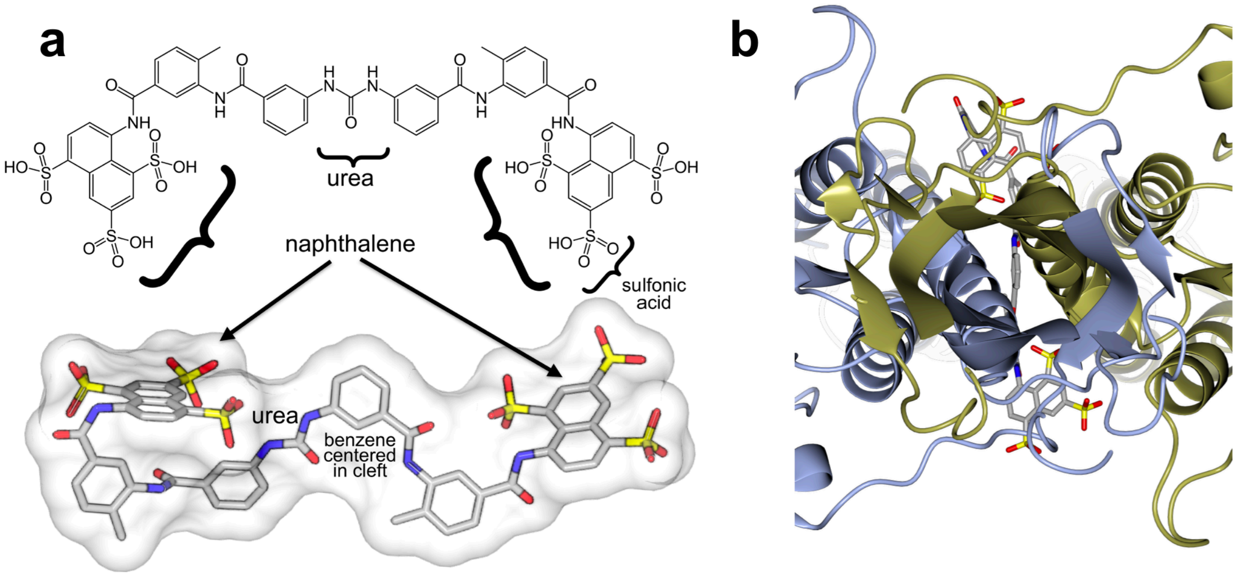


**Supplementary Fig. 6: Two-fold axis of dimer symmetry for VSGsur-suramin complex**.

**(a)** The top drawing illustrates the chemical structure of suramin with several of its functional groups denoted, whereas the bottom renders the drug as found in the protein structure with a transparent molecular surface shown. Oxygen atoms are shown in red, nitrogen in blue, and carbon gray. **(b)** Ribbon diagram of VSGsur (one monomer blue and the other gold) looking down the two-fold axis of symmrtry for the dimer. Suramin is shown as a ball-and-stick chemical representation as in (a). In the center of the rotational axis for the dimer, one of the suramin benzene rings is visible.

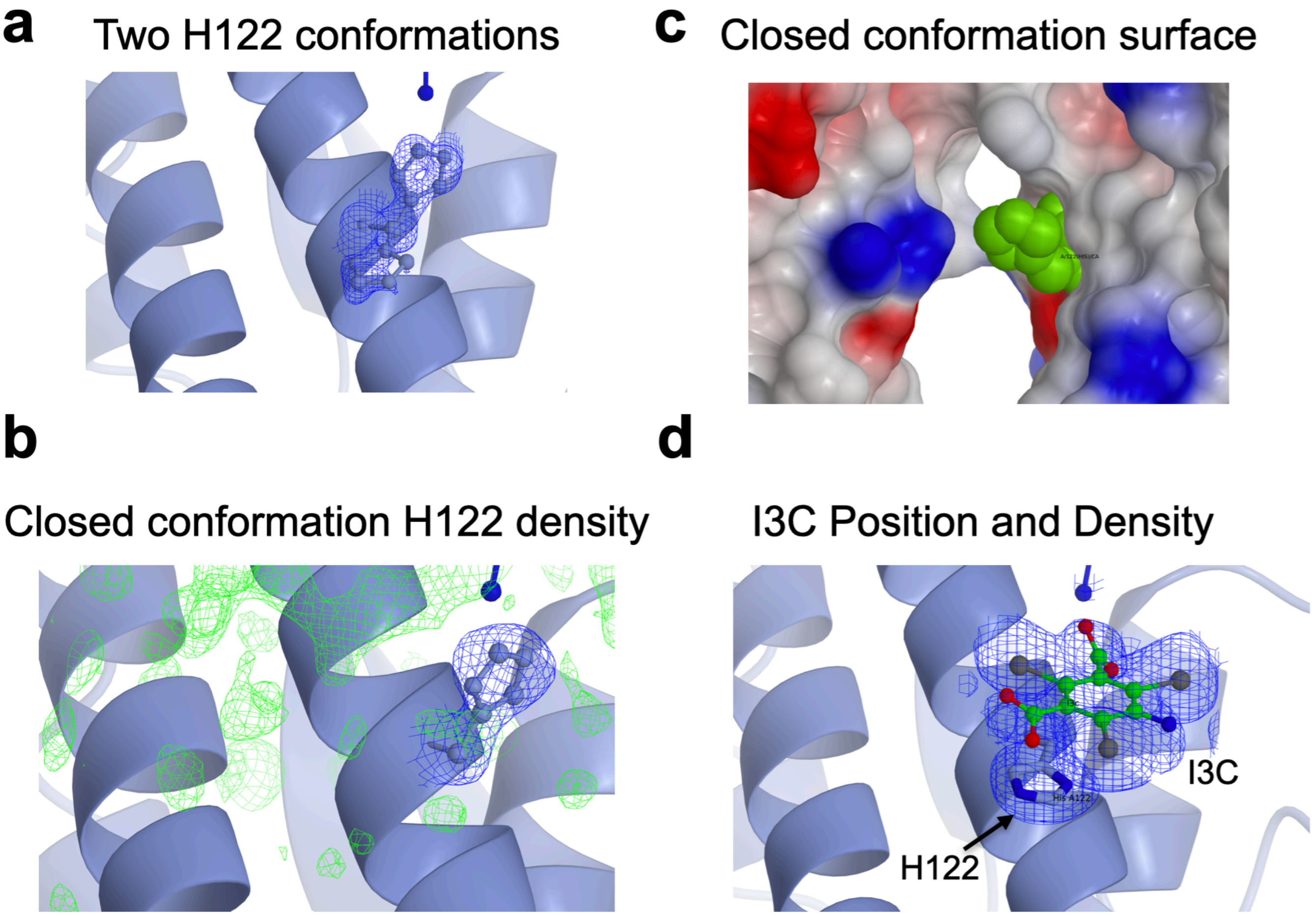


**Supplementary Fig. 7: H122 “open” and “closed” conformations.** (a) Two H122 conformations in the native crystals structure of VSGsur with corresponding electron density (b) Closed conformation electron density of H122 (c) Closed conformation surface of VSGsur (d) Position of H122 and the I3C group used in phasing the crystal structure with corresponding electron density.

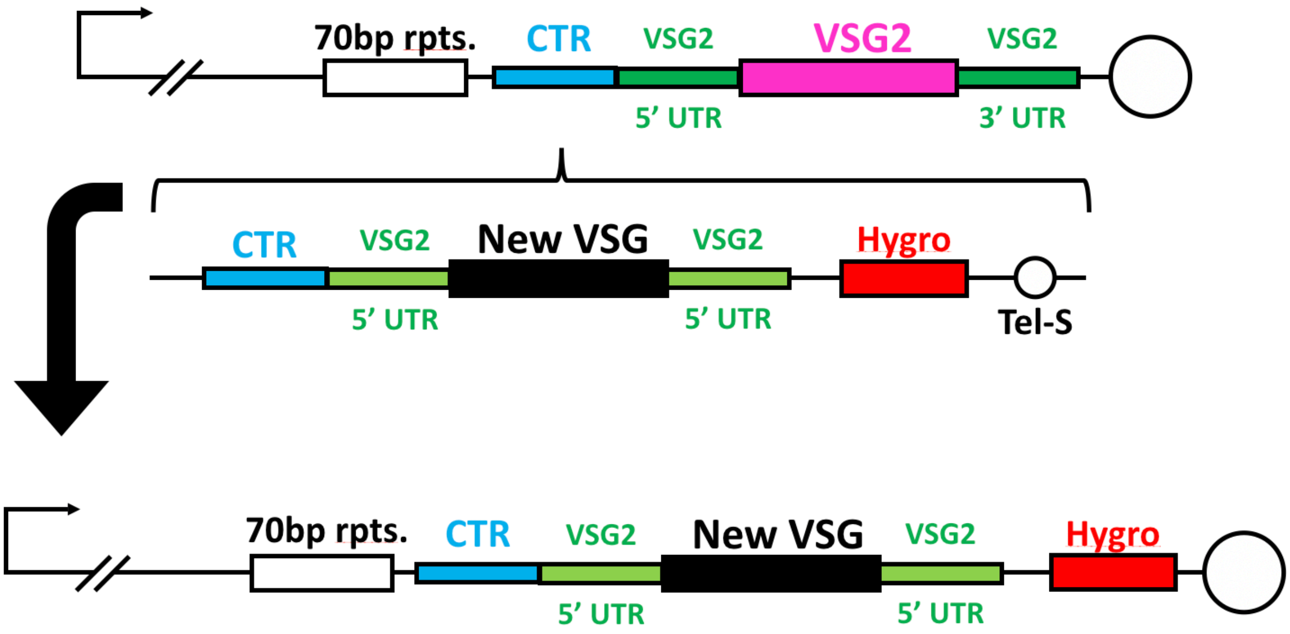


**Supplementary Fig. 8: Creation of VSGsur and VSGsur mutant expressing strains**
The same construct was used multiple times to generate different VSG-expressing cell lines. The top schematic shows the endogenous sub-telomeric expression site in 2T1 cells, which express VSG2 (pink) with its endogenous UTRs (dark green). The vector used to integrate new VSGs was adapted from Pinger et al.[1], wherein the homology directed integration of a novel VSG ORF (black) flanked by the UTRs of VSG2 is mediated by 3’ telomere seeds (Tel-S) and 5’ homology to the upstream co-transposed region (CTR – teal). Transfected cells are identified by screening for the integration of a hygromycin selection cassette (red) that is expressed via read-through transcription driven by the subtelomeric promoter.

### Panels of WT and H122A with suramin soaks

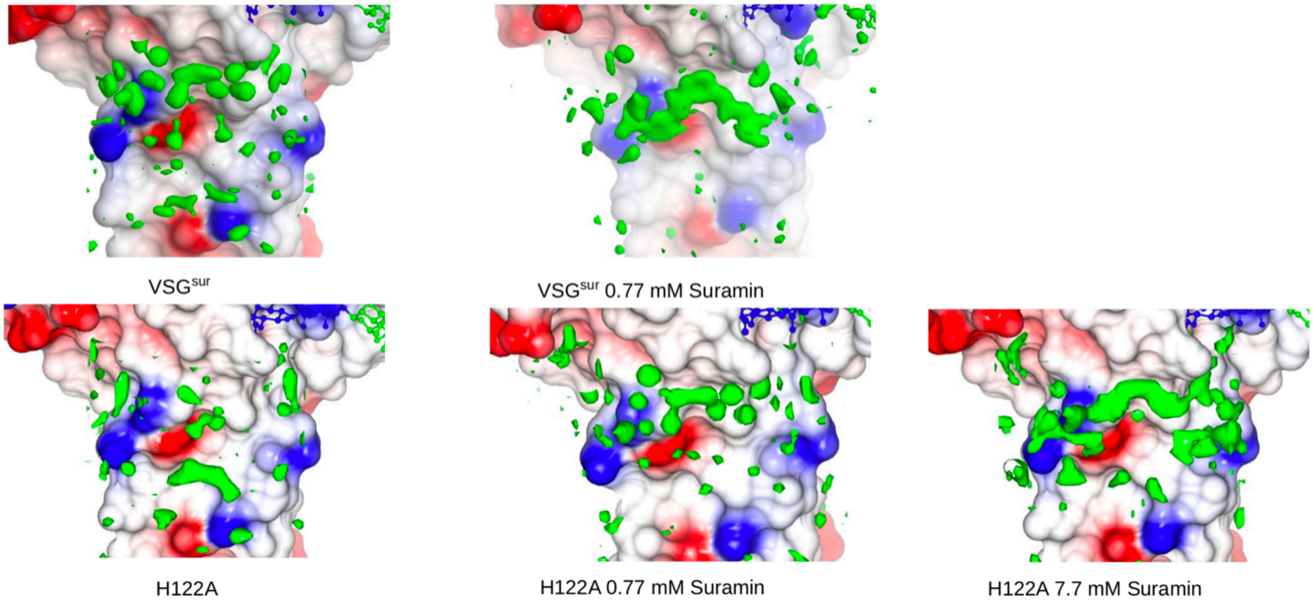


**Supplementry Fig. 9: Panels of WT and H122A with suramin soaks.** Molecular surface of one monomer in the VSGsur dimer shown colored by electrostatic potential (white is neutral, blue is positive/basic, and red is acidic/negatively charged). Electron density in the suramin binding site is shown in green.

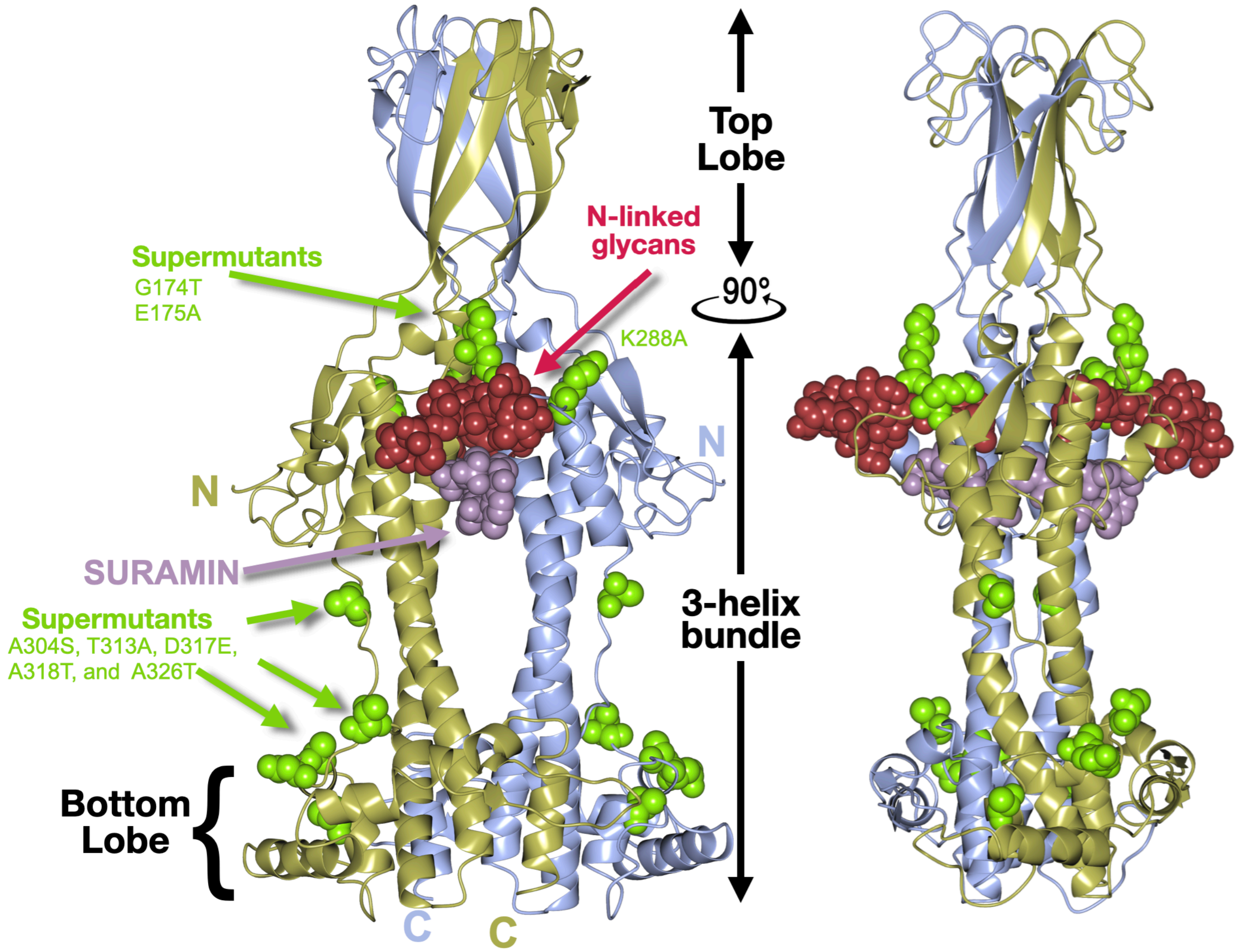


**Supplementary Fig. 10: "Supermutations" mapped to VSGsur.** Ribbon diagrams of VSGsur/suramin co-crystal structure in two orientations. Mutations discovered in "supersur" VSGsur mutants with heightened resistance to suramin are shown as green space filling atoms. Suramin and the N-linked glycan of VSGsur are depicted as space filling atoms in purple and crimson respectively.

a

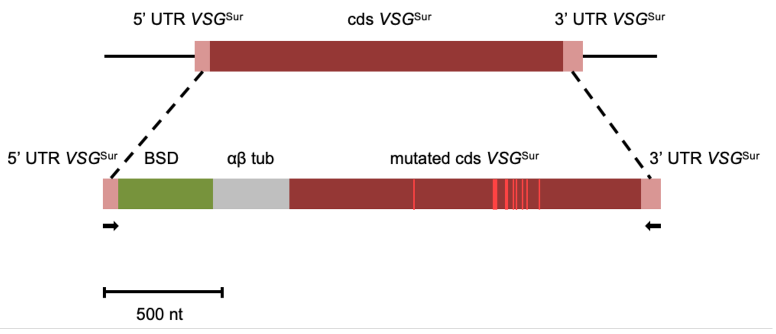


b

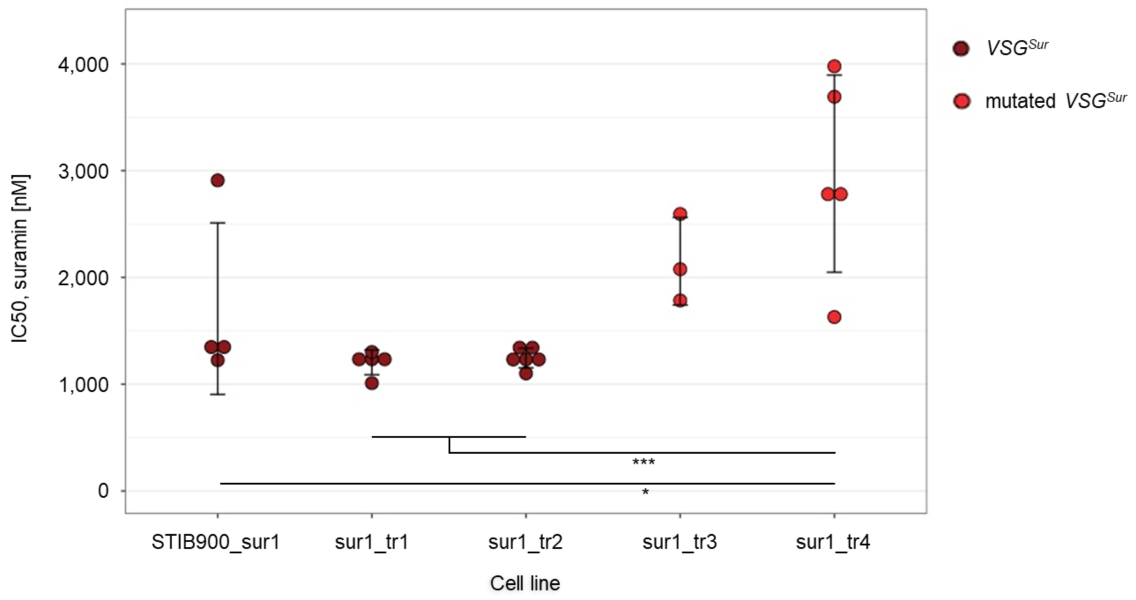


**Supplementary Figure 11. Replacement of VSGsur with "supermutant" VSGsur at the active expression site of *T. b. rhodesiense* STIB900_sur1.**

**(a)** Construct for gene replacement; mutations in the VSGsur coding sequence (dark red) are shown in light red, arrows indicate primer binding sites. BSD, blasticidine resistance gene; αβ tub, αβ tubulin splice site. **(b)** 50% inhibitory concentrations of the transfected clones and the parent (sur1) as measured with Alamar Blue assays. The pseudoclones sur1_tr1 and sur1_tr2 still expressed VSGsur, while the pseudoclones sur1_tr3 and sur1_tr4 expressed the mutant version. The scatter plots represent mean ± standard deviation. Significant differences are indicated (Tukey's multiple comparisons test).

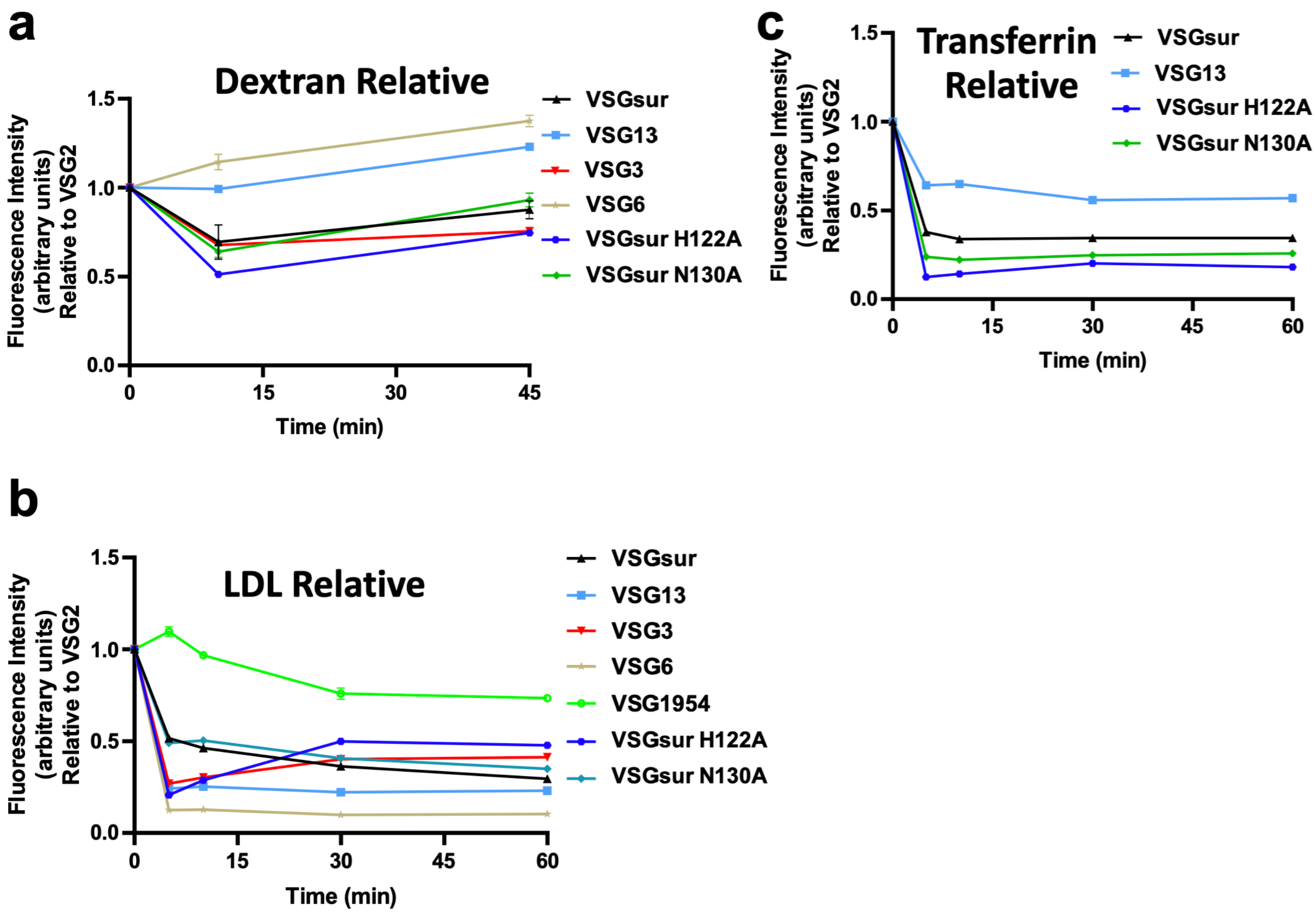


**Supplementary Fig. 12: Endocytosis rates by different VSGs relative to VSG2**

Alexa 488-dextran (a), bodipy-LDL (b), and Alexa 488-transferrin (c) endocytosis by *T. b. brucei* 2T1 cells expressing a variety of different VSG genes. Due to technical limitations, each cell line could not be analyzed simultaneously within each experiment for each ligand. Therefore, VSG2 was used as a control in all experiments, allowing the calculation of each cell line's relative uptake rate of each ligand as compared to the uptake rate of each ligand observed by VSG2 expressing cells within each separate experiment. Each of these graphs therefore represent the combined experimental results from 2 to 3 separate experiments. Cell lines with a relative fluorescence intensity below 1 at a given time point have less efficient endocytic rates compared to VSG2 expressing cells, and vice versa. All graphs share the same Y axis. Error bars when present denote the range (N=2 experimental replicates for each cell line at every time point), while error bars were left out when they were smaller than the size of the data point and therefore not visible.

#### Supplementary Table 1: Crystallographic Statistics for Wild Type VSGsur

| | VSGsur | VSGsur+I3C | VSGsur + 0.77mM suramin |
|---|---|---|---|
| **Data Collection** | | | |
| Beamline | BESSY MX 14.1 | BESSY MX 14.2 | SLS X06DA |
| Processing software | XDSAPP | XDSAPP | go.pi |
| Wavelength (Å) | 0.9184 | 2.066 | 1.0 |
| Resolution range (Å) | 48.05 -1.21 (1.25-1.21) | 44.15-1.92 (1.99-1.92) | 47.87-1.86 (1.93 -1.86) |
| Space group | P 21 21 2 | P 21 21 2 | P 1 21 1 |
| Unit Cell *a*, b, *c* (Å) | 47.0 71.0 130.4 | 46.96 70.89 129.65 | 52.69 79.22 118.00 |
| Unit Cell α, β, γ (°) | 90 90 90 | 90 90 90 | 90 90.78 90 |
| Total reflections | 747748 (26978) | 507768 (5366) | 545357 (55408) |
| Unique reflections | 122558 (7073) | 27999 (845) | 81516 (8077) |
| Multiplicity | 6.1 (3.8) | 18.1 (6.4) | 6.7 (6.8) |
| Completeness (%) | 91.49 (53.39) | 82.57 (25.54) | 99.91 (99.77) |
| Mean *I/sigma(I)* | 8.57 (0.38) | 13.99 (0.68) | 12.25 (2.32) |
| Wilson B-factor | 15.43 | 30.05 | |
| *R-merge* | 0.09566 (2.578) | 0.1762 (1.899) | 0.1091 (0.759) |
| *R-meas* | 0.1044 (2.988) | 0.1813 (2.055) | 0.1185 (0.8029) |
| *R-pim* | 0.04114 (1.487) | 0.04182 (0.7539) | 0.04537 (0.3039) |
| CC1/2 | 0.998 (0.174) | 0.995 (0.380) | 0.997 (0.741) |
| CC* | 0.999 (0.544) | 0.999 (0.742) | 0.999 (0.923) |
| **Refinement** | | | |
| Refinement reflections | 122545 (7073) | 27993 (845) | 81516 (8077) |
| R-free reflections | 2100 (121) | 1124 (34) | 3910 (372) |
| R-work | 0.174 (0.377) | 0.235 (0.333) | 0.195 (0.297) |
| R-free | 0.184 (0.348) | 0.256 (0.382) | 0.227 (0.337) |
| CC(work) | 0.964 (0.477) | 0.917 (0.588) | 0.910 (0.719) |
| CC(free) | 0.971 (0.494) | 0.933 (0.588) | 0.891 (0.597) |
| No. of atoms | 3305 | 2781 | 6654 |
| macromolecules | 2799 | 2481 | 5707 |
| ligands | 83 | 77 | 296 |
| solvent | 423 | 233 | 651 |
| Protein residues | 377 | 335 | 758 |
| RMS(bonds) | 0.01 | 0.008 | 0.008 |
| RMS(angles) | 1.41 | 1.04 | 0.94 |
| Ramachandran favored (%) | 98.36 | 98.46 | 97.33 |
| Ramachandran allowed (%) | 1.37 | 1.54 | 2.54 |
| Ramachandran outliers (%) | 0.27 | 0 | 0.13 |
| Rotamer outliers (%) | 0 | 0.80 | 0.84 |
| Clashscore | 3.3 | 1.60 | 5.54 |
| Average B-factor ($Å^2$) | 32.2 | 45.96 | 21.46 |
| macromolecules | 31.14 | 46.05 | 20.52 |
| ligands | 32.9 | 49.86 | 32.26 |
| solvent | 39.11 | 43.55 | 24.82 |
| Number of TLS groups | | 7 | 16 |

Highest-resolution shell statistics are in parentheses.

#### Supplementary Table 2: Crystallographic Statistics for VSG13

| | VSG13 | VSG13 + 0.5mM NaBr |
|---|---|---|
| **Data Collection** | | |
| Beamline | ESRF ID29 | Diamond i03 |
| Processing software | iMosflm /CCP4 DIALS | iMosflm/CCP4 DIALS |
| Wavelength (Å) | 1.0 | 0.9198 |
| Resolution range (Å) | 48.14 -1.378 (1.428- 1.378) | 52.55-1.56 (1.616-1.56) |
| Space group | C 1 2 1 | C 1 2 1 |
| Unit Cell *a*, b, *c* (Å) | 73.717 68.341 156.903 | 74.1477 68.4071 157.759 |
| Unit Cell α, β, γ (°) | 90 92.548 90 | 90 92.1646 90 |
| Total reflections | 465941 (45521) | 432679 (21591) |
| Unique reflections | 155980 (15141) | 109295 (8780) |
| Multiplicity | 3.0 (3.0) | 4.0 (2.4) |
| Completeness (%) | 97.09 (94.49) | 97.22 (78.32) |
| Mean *I/sigma(I)* | 8.07 (1.15) | 18.77 (0.09) |
| Wilson B-factor | 18.5 | 19.34 |
| *R-merge* | 0.04926 (0.6997) | 0.1892 (4.641) |
| *R-meas* | 0.05999 (0.8474) | 0.2143 (5.751) |
| *R-pim* | 0.03368 (0.4707) | 0.09891 (3.314) |
| CC1/2 | 0.995 (0.84) | 0.983 (0.06) |
| CC* | 0.999 (0.955) | 0.996 (0.336) |
| **Refinement** | | |
| Refinement reflections | 155980 (15092) | 109082 (8755) |
| R-free reflections | 1636 (159) | 2019 (164) |
| R-work | 0.2176 (0.4012) | 0.2263 (0.3773) |
| R-free | 0.2409 (0.4511) | 0.2325 (0.3552) |
| CC(work) | 0.949 (0.864) | 0.901 (0.233) |
| CC(free) | 0.947 (0.844) | 0.900 (0.200) |
| No. of atoms | 5782 | 5808 |
| macromolecules | 5256 | 5142 |
| ligands | 105 | 128 |
| solvent | 421 | 538 |
| Protein residues | 701 | 695 |
| RMS(bonds) | 0.011 | 0.009 |
| RMS(angles) | 1.12 | 0.95 |
| Ramachandran favored (%) | 97.67 | 98.09 |
| Ramachandran allowed (%) | 2.33 | 1.91 |
| Ramachandran outliers (%) | 0 | 0.00 |
| Rotamer outliers (%) | 0.71 | 0.00 |
| Clashscore | 2.81 | 4.23 |
| Average B-factor ($Å^2$) | 31.18 | 32.93 |
| macromolecules | 30.99 | 31.85 |
| ligands | 33.83 | 40.09 |
| solvent | 32.83 | 41.47 |
| Number of TLS groups | 0 | 28 |

Highest-resolution shell statistics are in parentheses.

#### Supplementary Table 3: Crystallographic Statistics for Mutant VSGsur

| | VSGsur H122A | VSGsur H122A + 0.77mM suramin | VSGsur H122A + 7.7mM suramin |
|---|---|---|---|
| **Data Collection** | | | |
| Beamline | Diamond i03 | SLS X06DA | SLS X06DA |
| Processing software | XIA2/DIALS (CCP4) | XIA2/DIALS (CCP4) | go.pi |
| Wavelength (Å) | 0.9763 | 1.0 | 1.0 |
| Resolution range (Å) | 48.05 -1.47 (1.523-1.47) | 38.08-1.75 (1.81-1.75) | 39.16-1.66 (1.72-1.66) |
| Space group | P 21 21 2 | P 21 21 2 | P 21 21 2 |
| Unit Cell *a*, b, *c* (Å) | 47.0158 70.9113 130.677 | 46.90 70.95 130.44 | 46.94 71.03 130.61 |
| Unit Cell $\alpha$, $\beta$, $\gamma$ (°) | 90 90 90 | 90 90 90 | 90 90 90 |
| Total reflections | 995504 (99575) | 588342 (59570) | 686746 (66206) |
| Unique reflections | 75170 (7360) | 44210 (4304) | 52136 (5028) |
| Multiplicity | 13.2 (13.5) | 13.3 (13.8) | 13.2 (13.2) |
| Completeness (%) | 99.28 (94.57) | 98.94 (97.44) | 99.79 (98.15) |
| Mean *I/sigma(I)* | 12.12 (0.44) | 16.67 (1.15) | 19.44 (1.49) |
| Wilson B-factor | 23.6 | 26.28 | 24.19 |
| *R-merge* | 0.09745 (2.95) | 0.1224 (2.281) | 0.0916 (1.719) |
| *R-meas* | 0.1014 (3.067) | 0.1273 (2.366) | 0.0953 (1.788) |
| *R-pim* | 0.02776 (0.8326) | 0.03457 (0.6261) | 0.0261 (0.4861) |
| CC1/2 | 1 (0.414) | 0.999 (0.503) | 1 (0.559) |
| CC* | 1 (0.766) | 1 (0.818) | 1 (0.847) |
| **Refinement** | | | |
| Refinement reflections | 74634 (6960) | 44206 (4304) | 52131 (5028) |
| R-free reflections | 3654 (349) | 2211 (215) | 2605 (251) |
| R-work | 0.2008 (0.3851) | 0.178 (0.325) | 0.185 (0.302) |
| R-free | 0.2236 (0.3853) | 0.207 (0.325) | 0.208 (0.320) |
| CC(work) | 0.959 (0.713) | 0.9579 (0.725) | 0.950 (0.787) |
| CC(free) | 0.946 (0.633) | 0.959 (0.726) | 0.950 (0.738) |
| No. of atoms | 3150 | 3067 | 2988 |
| macromolecules | 2749 | 2680 | 2644 |
| ligands | 83 | 72 | 61 |
| solvent | 318 | 315 | 283 |
| Protein residues | 368 | 359 | 356 |
| RMS(bonds) | 0.006 | 0.015 | 0.013 |
| RMS(angles) | 1.21 | 1.23 | 1.19 |
| Ramachandran favored (%) | 98.9 | 99.43 | 98.86 |
| Ramachandran allowed (%) | 1.1 | 0.57 | 1.14 |
| Ramachandran outliers (%) | 0 | 0 | 0 |
| Rotamer outliers (%) | 2.46 | 0.36 | 0 |
| Clashscore | 4.43 | 0.73 | 0.93 |
| Average B-factor ($Å^2$) | 38.44 | 39.22 | 38.91 |
| macromolecules | 38.26 | 38.90 | 38.86 |
| ligands | 39.85 | 36.81 | 30 |
| solvent | 39.6 | 42.47 | 41.31 |
| Number of TLS groups | 9 | 5 | |

Highest-resolution shell statistics are in parentheses.

#### SUPPLEMENTARY METHODS

##### 1. Construction of pHH-VSG$^{Sur}$-hyg plasmid:

A switched variant of VSG termed VSG$^{Sur}$ that emerged from *Trypanosoma brucei rhodesiense* culture under suramin selection has been reported to correlate with suramin resistance (Wiedemar *et al.*, Molecular Microbiology, 2018). A DNA sequence encoding the VSG$^{Sur}$ (GenBank: MF093647.1) was codon-optimized and synthesized as a pUC19 clone (BioCat, Germany). VSG$^{Sur}$ DNA was PCR amplified from pUC19-VSG$^{Sur}$ plasmid using Q5 DNA polymerase (New England Biolabs) and the following primers:

VSG$^{Sur}$-pHH-F1: CGACACGTACGCGGCATGCAAGCCGTAACACGC
VSG$^{Sur}$-pHH-R1: GAAATTTGAGGGGGGAAATTAAAAAAGCAAAAATGCAAGCAAAAGAGG

Also, a puromycin-resistant knock-in vector pHH-VSG3-PAC (PCT/EP2019/079063) was linearized by PCR using the following primers:

pHH-VSG2.3′-UTR-F1: TTTCCCCCCTCAAATTTCCCCCCTCC
pHH-VSG2-CTR-R1: ATGCCGCGTACGTGTCG

The VSG$^{Sur}$ and pHH knock-in vector amplicons were assembled using HiFi® DNA Assembly Master Mix (New England Biolabs) according to the manufacturer's instructions, to replace VSG3 with VSG$^{Sur}$. To replace puromycin resistance with hygromycin, a hygromycine gene was PCR amplified from pHD789 plasmid (Irmer and Clayton, Nucleic Acids Research, 2001) using the following primers:

Hyg-pHH-F1: GCTCTAGAACTAGTCAGCTTACCATGAAAAAGCCTGAACTCACCGCGAC
Hyg-pHH-R1: TGGGCAGGATCGATCCCTACTCTATTCCTTTGCCCTCGGACGAGTG

Also, the pHH knock-in vector excluding puromycin gene was linearized using the following primers:

Aldolase-3′-UTR-F2: GGATCGATCCTGCCCATTTGGCTTTTCCCTTGTCTCGTG
Actin-5′-UTR-R2: AGCTGACTAGTTCTAGAGCTTATTTTATGGCAGCAACGAGACCTTAC

Finally, the hyg gene and pHH knock-in vector amplicons were assembled using HiFi® DNA Assembly Master Mix (New England Biolabs) according to the manufacturer's instructions.

##### 2. Generation of a *T. brucei* Lister427 Clone Expressing VSG$^{Sur}$:

A *T. brucei brucei* cell line expressing VSG2, termed 2T1 (Alsford and Horn, Molecular Biochemical Parasitology, 2008), was transfected with EcoRV-linearized pHH-VSG$^{Sur}$-hyg plasmid to replace VSG2. Briefly, $2x10^7$ cells were electroporated with 10 µg DNA in 100 µl of a home-made Tb-BSF buffer (Burkard *et al.*, Molecular and Biochemical Parasitology, 2011) using Amaxa Nucleofector (Lonza), program X-001. After incubation in non-selective HMI-9 medium for 6 hours, hygromycin was added at 5 µg/ml and the cells were grown for 6 days at 37°C, 5% $CO_2$. Single clones isolated by serial dilution were screened for a VSG2-negative phenotype by FACS analysis using an APC-conjugated anti-VSG2 mouse mAb.

**3. Generation of pHH-VSG vector for VSG Knock-in *T. brucei*:**

Genomic DNA was extracted from VSG2-expressing 2T1 cells using DNAzol® reagent (Life Technologies) according to the manufacture's instruction. The VSG2 gene plus its upstream co-transposed region (CTR) and downstream telomere region were PCR amplified using Q5 DNA polymerase (New England Biolabs) and the following primers:

```
VSG2-CTR-F2:  GAAGGCAGCGGAAAGTGTGCCAATGC
Tb427-tel-R2: AACACCTTAATCCGAAACACC
```

Also, pUC19 vector (Life Technologies) was linearized by PCR using the following primers:

```
pUC19-tel-F1: GATATCTCTAGAGTCGACCTGCAGGCATGCAAGC
pUC19-CTR-R2: ACTTTCCGCTGCCTTCGATATCGGATCCCCGGGTACCG
```

Additionally, telomere seeds were amplified by PCR from pSY-37F1D-CTR-BSD plasmid (Pinger *et al*., Nature Communications, 2017) using the following primers:

```
pSY.tel-F1: ACGGTGTTTCGGATTAAGGCCGCGGGAATTCGATTAGG
pSY.tel-R1: AGGTCGACTCTAGAGATATCGGATCCACTAGCTAGTGATTAAC
```

Finally, the above mentioned amplicons were assembled to make pHH-VSG2-Tel plasmid using HiFi® DNA Assembly Master Mix (New England Biolabs) following the manufacturer's instructions. In order to insert a puromycin resistance cassette, pHH-VSG2-Tel was linearized by PCR using the following primers:

pHH-VSG2-Tel-F1:
```
ATGGGGGGATATTAGACTTAGGCTTAGGATTAGGATTAGGATTAGGATTAGGGTTAATTTTTTCCTCTTT
TTTTTTAACTCACACCTCTATCCTG
```

```
pHH-VSG2-Tel-R1: GCTTGCATGCCGCGTTCGTG
```

Also, a puromycin resistance cassette was PCR amplified using the following primers:

```
pHH-PAC-F1: GGATTAGGCACAGCAAGGTCTTCTGAAATTCATGT
pHH-PAC-R1: CTAAGTCTAATATCCCCCCATTTTCTTCTTTTACATCA
```

The PCR amplicons from pHH-VSG2-Tel vector, and the puromycin cassette were assembled to make pHH-VSG2-PAC plasmid using HiFi® DNA Assembly Master Mix (New England Biolabs) following the manufacturer's instructions. To replace puromycin resistance with hygromycin, a hygromycine gene was

PCR amplified from pHD789 plasmid (Irmer and Clayton, Nucleic Acids Research, 2001) using the following primers:

```
Hyg-pHH-F1: GCTCTAGAACTAGTCAGCTTACCATGAAAAAGCCTGAACTCACCGCGAC
Hyg-pHH-R1: TGGGCAGGATCGATCCCTACTCTATTCCTTTGCCCTCGGACGAGTG
```

Also, the pHH knock-in vector excluding puromycin gene was linearized using the following primers:

```
Aldolase-3′-UTR-F2: GGATCGATCCTGCCCATTTGGCTTTTCCCTTGTCTCGTG
Actin-5′-UTR-R2: AGCTGACTAGTTCTAGAGCTTATTTTATGGCAGCAACGAGACCTTAC
```

Finally, the hyg gene and pHH knock-in vector amplicons were assembled using HiFi® DNA Assembly Master Mix (New England Biolabs) according to the manufacturer's instructions.

**Alamar Blue assays**

Serial drug dilutions were prepared on a 96-well plate and the parasites were added to a final concentration of $2 \times 10^4$ cells/ml. After incubation for 68 hours, resazurin was added to a concentration of 11.4 µg/ml. The fluorescence of viable cells was determined after 3-4 hours with a SpectraMax reader (Molecular Devices) and SoftMax Pro 5.4.5 Software. Fitting of the dose-response curves (non-linear regression model, variable slope; four parameters, lowest value set to zero) and calculation of the IC50 values were carried out with GraphPad Prism 6.00.
